# Optoribogenetic control of regulatory RNA molecules

**DOI:** 10.1101/2020.07.07.191379

**Authors:** Sebastian Pilsl, Charles Morgan, Moujab Choukeife, Andreas Möglich, Günter Mayer

## Abstract

Short regulatory RNA molecules underpin gene expression and govern cellular state and physiology. To establish a novel layer of control over these processes, we generated chimeric regulatory RNAs that interact reversibly and light-dependently with the light-oxygen-voltage photoreceptor PAL. By harnessing this interaction, the function of micro RNAs (miRs) and short hairpin (sh) RNAs in mammalian cells can be regulated in spatiotemporally precise manner. The underlying strategy is generic and can be adapted to near-arbitrary target sequences. Owing to full genetic encodability, it establishes unprecedented optoribogenetic control of cell state and physiology. The method stands to facilitate the non-invasive, reversible and spatiotemporally resolved study of regulatory RNAs and protein function in cellular and organismal environments.

## Introduction

Short regulatory RNA molecules such as endogenous micro RNAs (miR) or synthetic short hairpin RNAs (shRNA) are essential mediators of gene expression^1-3^. They interact with defined complementary sites in the untranslated (UTR) or the coding regions of mRNA molecules, upon which translation is either inhibited or the mRNA is hydrolysed. Regulatory RNAs have become indispensable in the biosciences for the validation of gene or protein function in cells and *in vivo* ^4^. Although the on-demand control of mRNA translation has been achieved at the levels of mRNA stability and ribosome processing, e.g., by introducing aptazymes or aptamers in the UTRs^5-9^, the direct control of the function of short regulatory RNAs, ultimately in a spatiotemporal manner, remains challenging. At the same time, it is highly demanded, as it would offer programmable, modular and generalizable control of target gene expression on the posttranscriptional level^10-12^. To this end, small-molecule-responsive siRNAs whose function can be controlled by theophylline or tetracycline^13^ or conditional expressions systems of shRNAs^14^ have been reported. These approaches extend towards the transcriptional regulation of miRs^15^ or to aptazymes that control miR maturation in response to small molecules^16^. Atanasov *et al*. constructed *pre*-miR variants that functionally depend on the presence of doxycycline, mediated by a TetR-responsive aptamer^17^, which has been previously used in combination with the theophylline aptamer to control transcription^18^. Besides these strategies, modalities to sequester miRs^19-21^ or to inhibit their function by small molecules^22^ were developed and applied in cell culture and *in vivo*. Most of these approaches rely on the exogenous addition of small molecules, which *per se* might interfere with other biological processes, have limited availability and stability *in vivo*, suffer from diffusional spread, and are of restricted reversibility^23^. To overcome certain of these limitations, light-dependent control of regulatory RNA has also been described^24-26^, but the pertinent approaches invariably require chemical synthesis and the exogenous addition of the modified RNAs to biological systems. By contrast, entirely genetic approaches to gain spatiotemporal control over regulatory RNA function remain elusive but are highly desirable, as they would offer a plethora of applications to precisely and reversibly control gene expression and downstream processes.

Here, we devise a fully genetically encodable, generic approach that achieves light-dependent control of *pre*-miR and shRNA activity. We constructed chimeric RNA molecules consisting of mature miR and siRNA sequences conjoined with an RNA aptamer that binds to the light-oxygen-voltage (LOV) photoreceptor PAL in a light-dependent manner^27,28^. The chimeric RNAs enable the spatiotemporal control of short regulatory RNA function in mammalian cells, as we showcase for the light-dependent control of gene expression and cell-cycle progression. This hitherto unavailable modality establishes a versatile RNA control system for analysing various protein and miR functionalities in a reversible, spatiotemporally resolved, and non-invasive manner, and with full genetic encoding. Owing to the modularity of the chimeric RNAs, the technology readily applies to near-arbitrary shRNAs.

## Results

### PAL-mediated regulation of *pre*-miR activity

Our design of light-responsive *pre*-miRs anticipates an altered processivity of short regulatory RNAs by Dicer owing to light-activated binding of the light-oxygen-voltage receptor PAL to the apical loop domain^28-30^. To implement this design, we embedded the cognate aptamer domain of PAL in the apical loop of short regulatory RNAs. We hypothesized that thereby regulatory RNA function can be controlled by blue light (**Scheme 1**).

**Scheme 1:**
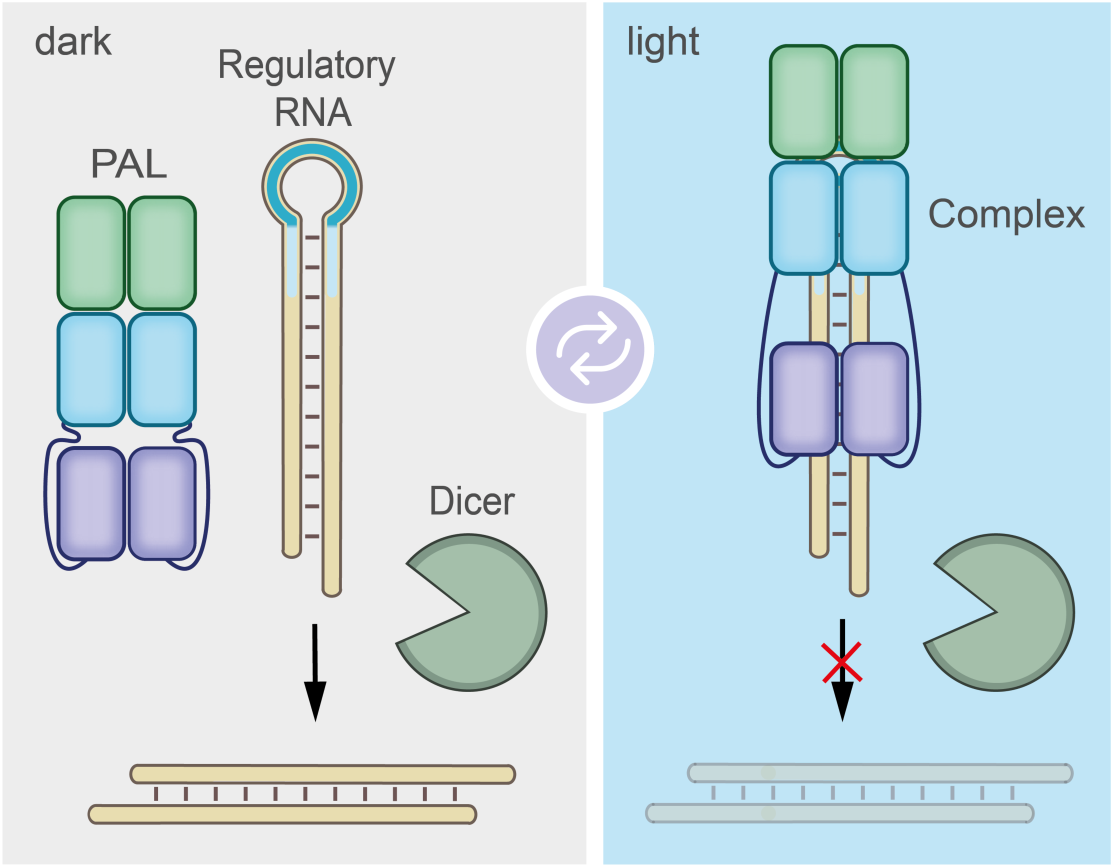
General design of light-dependent regulatory RNAs. The PAL protein reversibly binds to its cognate RNA aptamer (highlighted in blue) embedded in the apical loop domain of a regulatory RNA (highlighted in light orange) in the light and thereby influences regulatory RNA function.

We generated *pre*-miR variants by replacing the apical loop domain with the PAL binding RNA aptamer 53 (**Fig. 1a, Supporting Table 1**). Notably, the RNA aptamer 53 interacts preferentially with the light-adapted state of PAL and to a much lesser extent with its dark-adapted conformation (**Supporting Fig. 3a,b**). We first generated aptamer-modified variants of *pre*-miR-21 (SHA, **Fig. 1a**) and analysed them in reporter gene assays that employ the expression of secreted *Metrida* luciferase or of enhanced green fluorescent protein (eGFP) with miR-21 target sites embedded in the 3’-UTRs of the respective mRNA (**Supporting Fig. 1a,b,2**)^31^. As controls, we constructed *pre*-miR-21 variants that bear a single point mutant (G11C) within the PAL aptamer that renders them binding-incompetent (SHC, SHD), a non-functional miR-21 domain (SHB, SHD) ^32^, or with both domains altered (SHD, **Fig. 1a**). Interaction experiments *in vitro* revealed light-dependent binding of SHA and SHB to PAL, similar to the parental aptamer (53), whereas *pre*-miR variants with mutated aptamer domains (SHC, SHD) did not bind (**Supporting Fig. 3a,b**). For all experiments, a transgenic HEK293 cell line stably expressing mCherry-PAL (HEK293PAL) at an average concentration of 1 µM was used (**Supporting Fig. 4**). The *pre*-miR-21 variants were transcribed under the control of the U6 promoter from plasmids^33^ co-transfected with the luciferase reporter. Whereas SHA supressed luciferase expression in darkness, irradiation with blue light (λ = 465 nm) induced reporter gene expression by 4.4-fold to 27% of the maximal value (**Fig. 1b,d**). Replacing either the miR-21 domain with a non-targeting RNA (SHB) or the aptamer domain by a non-binding point mutant (SHC) resulted in a loss of light-regulation (**Fig. 1b,d, Supporting Fig. 5**). Likewise, the *pre*-miR21 variant having both RNA domains altered (SHD) neither supressed gene expression nor showed any light dependency (**Fig. 1b,c**). Analogous results were obtained for the eGFP reporter gene (**Fig. 1d,e, Supporting Fig. 7,8**, for details on eGFP gating strategy see **Supporting Fig. 6a**), in that SHA inhibited expression in darkness, whereas a 4.4-fold induction of eGFP was observed in light (**Fig. 1d,e**). SHB and SHD did not inhibit eGFP expression, whereas SHC did, and none of the three variants exhibited light dependency (**Fig. 1d**). Intrinsic levels of argonaute 2 (AGO2) have been shown to limit RNA silencing efficiency^34^. Therefore, we co-expressed AGO2 and observed a more pronounced inhibition of eGFP expression by SHA and SHC in darkness (**Fig. 1f,g, Supporting Fig. 9,10**). Irradiation induced eGFP expression in the cells harboring SHA by 9-fold (**Fig. 1g**). By contrast, experiments using SHB, SHC and SHD did not reveal any light dependency (**Fig. 1f,g**).

**Fig. 1.**
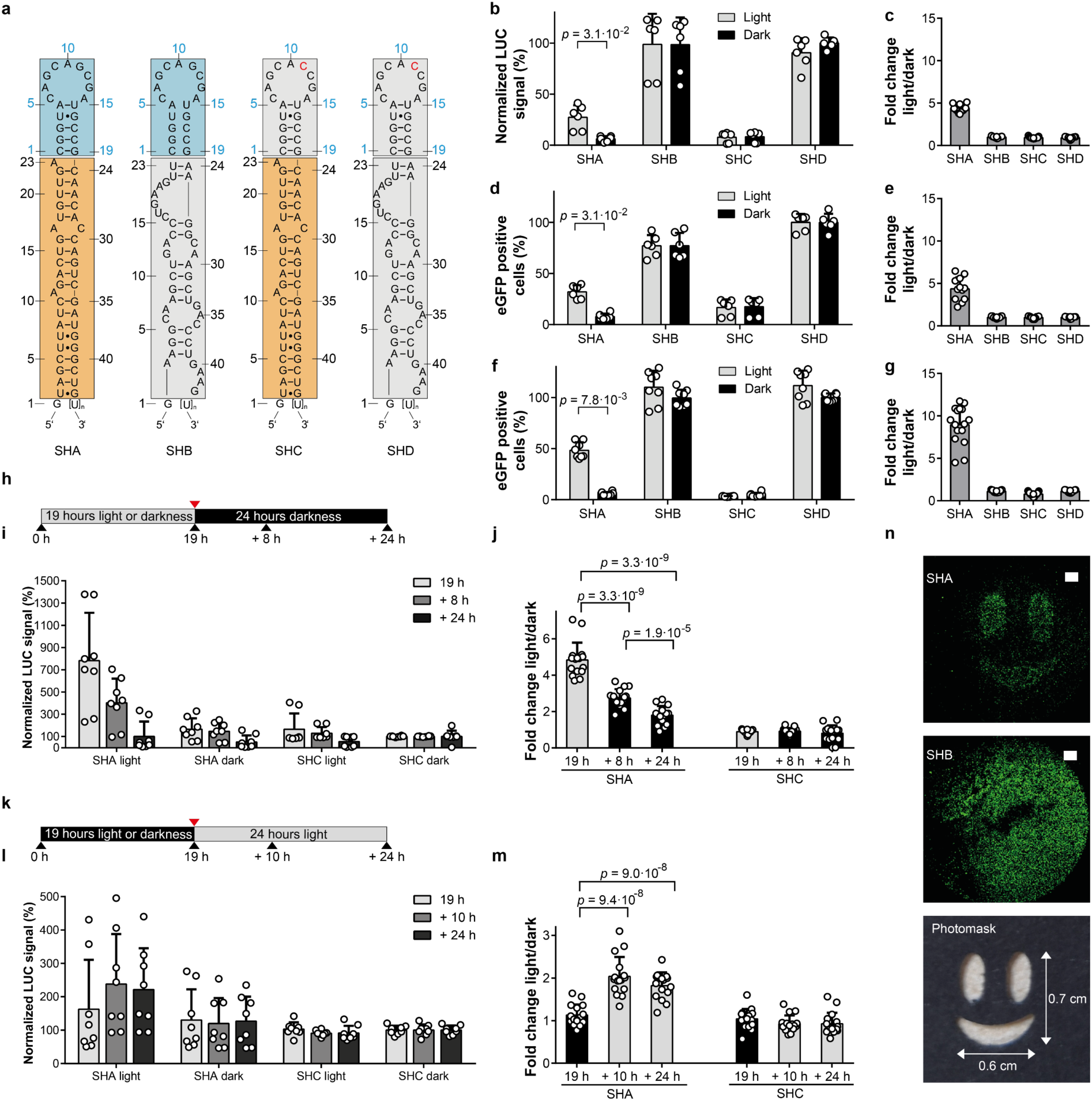
A *pre*-miR21-aptamer chimera enables light-control of gene expression. **a**, Schematic representation of the *pre*-miR21 variants and corresponding controls. Blue boxes: aptamer domain, orange boxes: miR21 domain, grey boxes: aptamer point mutant or control miR. **b**, Luciferase expression after transfection of the indicated *pre*-miR21 variants. Values are normalized to SHD incubated in darkness. **c**, Fold changes calculated from light *vs*. dark conditions from (**b**). **d**, Number of cells expressing eGFP after transfection of the indicated *pre-*miR21 variants. Values are normalized to SHD incubated in darkness. **e**, Fold changes calculated from light *vs*. dark conditions from (**d**). **f**, Number of cells expressing eGFP in the presence of elevated levels of AGO2 and after transfection of the indicated *pre*-miR21 variants. Values are normalized to SHD incubated in darkness. **g**, Fold changes calculated from light *vs*. dark conditions from (**f**). **b-e**, N = three biologically independent experiments performed in duplicates. Grey bars: light conditions, black bars: dark conditions. Dark grey bars: fold changes. **c-g**, Grey bars: cells incubated under light conditions, black bars: cells incubated in darkness. **b-f**, Wilcoxon two-sided signed-rank test was used for statistical analysis as a paired observation was assumed. **h**, Illumination protocol applied in (**i**) and (**j**). **i**, Luciferase expression level of cells expressing SHA or SHC. Shown are normalized values to SHC in darkness. **j**, Fold changes calculated from light *vs*. dark conditions from (**i**). **k**, Illumination protocol applied in (**l**) and (**m**). **l**, Expression level of luciferase of cells expressing SHA or SHC. Shown are normalized values to SHC in darkness. **m**, Fold changes calculated from light *vs*. dark conditions from (**l**). **b-m**, Values are means ± s. d. **f-m** of four biologically independent cultures in duplicates. **j-m**, Two-sided Mann-Whitney *U* test was used for statistical analysis as an unpaired observation was assumed. **n**, Spatial patterning of eGFP expression after transfection with SHA (top panel) or SHB (middle panel). Irradiation was done on cells covered with a photomask (bottom panel); white bars: 1000 µm.

We next assessed the reversibility of the approach using the luciferase reporter system. To this end, HEK293PAL cells harboring SHA were incubated for 19 h under blue light (**Fig. 1h-j, Supporting Fig. 11**). Subsequently, the cells were kept in darkness for a further 24 h. An increase of luciferase activity in the cell culture supernatants was observed after 19 h in light and a reduction when cells were kept in the dark afterwards (**Fig. 1i,j**). In turn, cells kept first in darkness did not reveal luciferase expression (**Fig. 1k-m, Supporting Fig. 12**), but luciferase activity was detected when cells were subsequently exposed to light conditions (**Fig. 1l,m**). Cells having SHC did not reveal light-dependent luciferase expression (**Fig. 1i,j,l,m, Supporting Fig. 11,12**). We also demonstrated spatial control of reporter gene expression using a photomask on HEK293PAL cells during irradiation (**Fig. 1n**). Expression of SHA resulted in eGFP expression predominantly in light-exposed areas, whereas eGFP expression was observed independently of the irradiation status in the presence of SHB (**Fig. 1n**).

To better characterize the processed miRs, we analysed them by 3’ miR-RACE (rapid amplification of cDNA ends). Compared to reported natural *pre*-miR-21, we observed altered processing of SHA at the 3’end of miR-21-5p (**Supporting Table 2**). We attribute this observation to using the U6 promotor for *pre*-miR-21 expression which requires an additional G-nucleotide for efficient transcription and, thus, induces altered Dicer processing^35^.

### PAL-mediated regulation of shRNA activity

We next investigated whether the PAL-aptamer system can also be applied to shRNA molecules in a more generic manner to thereby enable versatile optogenetic control of RNA interference^36^. Initially, we constructed two shRNAs (SH1, SH2) that target different sites within the eGFP mRNA coding region (**Supporting Fig. 1c**) and conjoined them with the PAL aptamer (**Fig. 2a,b**). The expression of eGFP in HEK293PAL cells harboring SH1 or SH2 was light-responsive, with SH1 being more efficient in eGFP suppression in the dark (**Fig. 2c,d, Supporting Fig. 13-16**). As a control we used the miR-21-targeting SHA (**Fig. 1a**), which did not inhibit eGFP expression (**Fig. 2c,d, Supporting Fig. 13-17**, for details on eGFP gating strategy see **Supporting Fig. 6b**), as the miR target site is absent in the reporter mRNA employed in this experiment (**Supporting Fig. 1c**). Structural variations of one or two nucleotides surrounding the Dicer cleavage site are common motifs found in natural *pre*-miRs and shRNAs^37^. These motifs alter the accuracy of shRNA processing and, thus, gene silencing efficacy^38^. We hence extended our study towards examining the impact of the nucleotides’ identity in the hinge region that connects the siRNA with the aptamer domain on shRNA performance. To this end, we designed 8 variants with single nucleotide bulges in the hinge region of SH1 (**Fig. 2d**) located either up-(SH3, SH4, SH5, SH6) or downstream (SH7, SH8, SH9, SH10) of the aptamer domain (**Fig. 2e**). All variants demonstrated light-dependent induction of eGFP expression but with varying efficiency (**Fig. 2f**). An upstream C (SH6) or a downstream G (SH9) nucleotide, relative to the aptamer domain, revealed the lowest expression, upstream A (SH3) or G (SH5) nucleotides or downstream A (SH7), U (SH8) or C (SH10) nucleotides exhibited very similar properties. Likewise, the suppression efficiency of shRNAs in the dark varied among the constructs (**Supporting Fig. 13-15**). The observed fold changes of eGFP expression (light *vs*. dark) are comparable across all shRNAs with SH9 having the lowest induction rate (**Fig. 2g**). In turn, an upstream U nucleotide (SH4) revealed similar fold changes (**Fig. 2g**) but a higher level of light-induced eGFP expression (**Fig. 2f**). Therefore, we chose A and U residues as representatives in the SH2 hinge region variants and included G (SH15) as a less efficient control. Single nucleotides inserted into the hinge regions of SH2 led to an improved eGFP knockdown in the dark (**Fig. 2h,i, Supporting Fig. 15**), and all variants remained light-responsive. An adenine (SH11) or uridine (SH12) nucleotide at the hinge region upstream of the aptamer domain led to the highest number of eGFP-positive cells (**Fig. 2i**), and SH14 revealed the strongest increase of eGFP expression upon irradiation (15.3. fold) (**Fig. 2j**). SH16 with a mutated aptamer domain (G11C) did not reveal light induced eGFP expression (**Fig. 2h,i**). Likewise, light-dependent induction of eGFP expression was also evident from fluorescence microscopy studies for on the shRNA variants (**Fig. 2k, Supporting Fig. 17**) SH1 (original hinge region), SH3 (intermediate performance), SH4 (highest number of eGFP positive cells when incubated in light), SH12 (second highest number of eGFP positive cells when incubated in light), and SH14 (highest light *vs*. dark fold change). *In vitro* binding studies verified light-dependent interaction with PAL of shRNA variants with engineered hinge regions (**Supporting Fig. 3a,b**), indicating that these variations do not directly interfere with PAL binding but affect shRNA processivity. Of key importance, these findings testify to the modular design of the underlying chimeric RNAs and indicate that the domains for PAL-binding and mRNA-targeting are non-overlapping. As a corollary, we reasoned that near-arbitrary targeting domains should be accommodable with our technology.

**Fig. 2.**
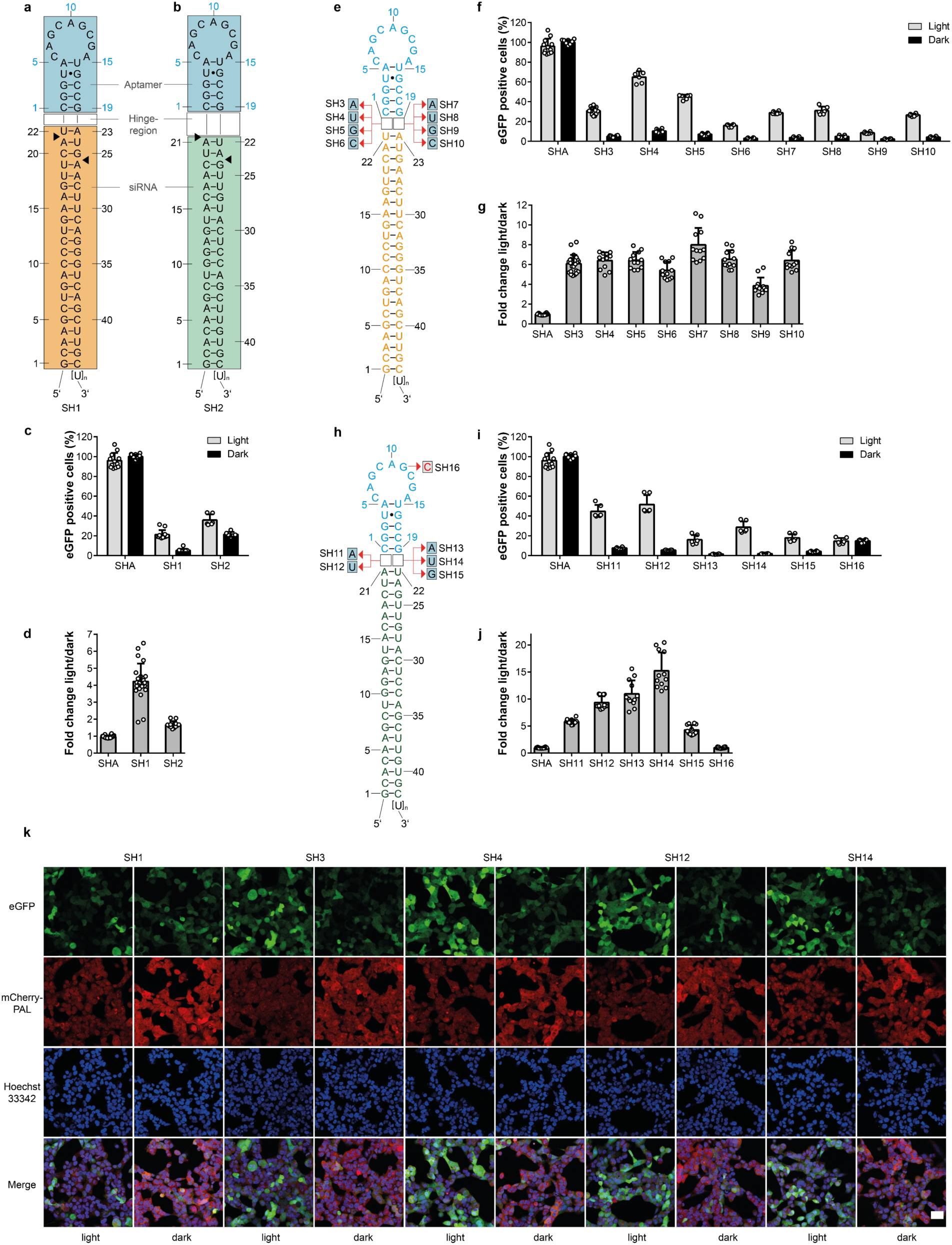
Design of shRNAs for the light-dependent expression of eGFP. Two different siRNA sequences SH1 (orange box, **a**) and SH2 (green box, **b**) targeting eGFP mRNA were conjoined with the PAL aptamer (blue boxes) as apical loop domains. Black arrows indicate a putative preferential dicer cleavage site^39^. **c**, Number of cells expressing eGFP after transfection of SH1 or SH2. Values are normalized to SHA (**Fig. 1a**) in darkness. **d**, Fold changes calculated from light *vs*. dark conditions from (**c**). **e**, Single nucleotide permutations of the hinge region in SH1 and their impact on eGFP expression and light-dependency (**f**). Values are normalized to SHA in darkness. **g**, Fold changes calculated from light *vs*. dark conditions from (**f**). **h**, Single nucleotide permutations of the hinge region in SH2 and their impact on eGFP expression and light-dependency (**i**). Values are normalized to SHA in darkness. **j**, Fold changes calculated from light *vs*. dark conditions from (**i**). **k**, Fluorescence microscopy images of cells transfected with the indicated shRNA variants. Cells were incubated under either light or dark conditions. Scale bar: 40 µm. **c-k**, All experiments were performed in duplicates and three independent replicates. Grey bars: light conditions, black bars: dark conditions. Dark grey bars: fold changes. Values are means ± s.d.

### Optoribogenetic control of cell cycle progression

We hence extended our approach to regulating the expression of endogenous proteins via shRNAs. We chose cyclin B1 and CDK1 as targets, as they are both essential for the transition from the gap-2 (G_2_) to the mitosis (M) phase of the cell cycle^40^. Variations of the expression levels of cyclin B1 and CDK1 have phenotypic consequences and alter the distribution of cells in different stages of the cell cycle^41,42^. First, we generated shRNAs targeting cyclin B1 with varied hinge nucleotides, having either an adenine (SHCB1) or uridine (SHCB2) upstream or uridine (SHCB3) downstream of the aptamer domain (**Fig. 3a**). HEK293PAL cells having the shRNAs SHCB1-3 in darkness (i.e. cyclin B1 knockdown condition) accumulated in the G_2_/M phase (**Fig. 3b, Supporting Fig. 18,19**). Upon irradiation, the number of cells in G_2_/M phase was significantly reduced in cells having SHCB1 (**Fig. 3b, Supporting Fig. 18,19**), indicating the recovery of normal cell cycle propagation. By contrast, propagation was not recovered upon irradiation for the binding-incompetent aptamer variants of SHCB1 (G11C, SHCB1m) (**Fig. 3b**). SHCB2 and SHCB3 did not affect cell cycle propagation when irradiated (**Fig. 3b**). Based on these results, we constructed PAL-dependent shRNA variants of CDK1 having an adenine nucleotide in the hinge region upstream of the aptamer (SHCDK1, **Fig. 3c**). HEK293PAL cells having the shRNAs SHCDK1 in darkness also accumulated in the G_2_/M phase (**Fig. 3d, Supporting Fig. 18,19**). Upon irradiation, the number of cells in G_2_/M phase was significantly reduced (**Fig. 3d, Supporting Fig. 18,19**). Cells having the PAL-binding deficient mutant shRNA SHCDK1m accumulated in the G_2_/M phase irrespective of the irradiation status (**Fig. 3b,d, Supporting Fig. 18,19**). No accumulation of cells in the G_2_/M phase was observed when cells expressed the non-targeting SH3 or were untreated (**Supporting Fig. 20**). However, a slight accumulation of cells in G_2_/M phase was observed upon irradiation (**Supporting Fig. 20**), most likely because of secondary irradiation effects on cells^43^. SHCB1 or SHCDK1 led to a decrease of cyclin B1 and CDK1 expression, respectively, which was reversed by irradiation (**Fig. 3e**-**g, Supporting Fig. 21**). Variants of the shRNAs deficient for PAL binding (SHCB1m, SHCDK1m) suppressed protein expression independently of light (**Fig. 3e**-**g**). The non-targeting shRNA SH3 (**Fig. 2b**) did not affect cyclin B1 and CDK1 expression, the expression levels of both proteins were similar those when cells were untreated (**Fig. 3e**-**g, Supporting Fig. 21**). Of note, the shRNA variants targeting cyclin B1 did not affect CDK1 expression and *vice versa* (**Fig. 3e, Supporting Fig. 21**).

**Fig. 3.**
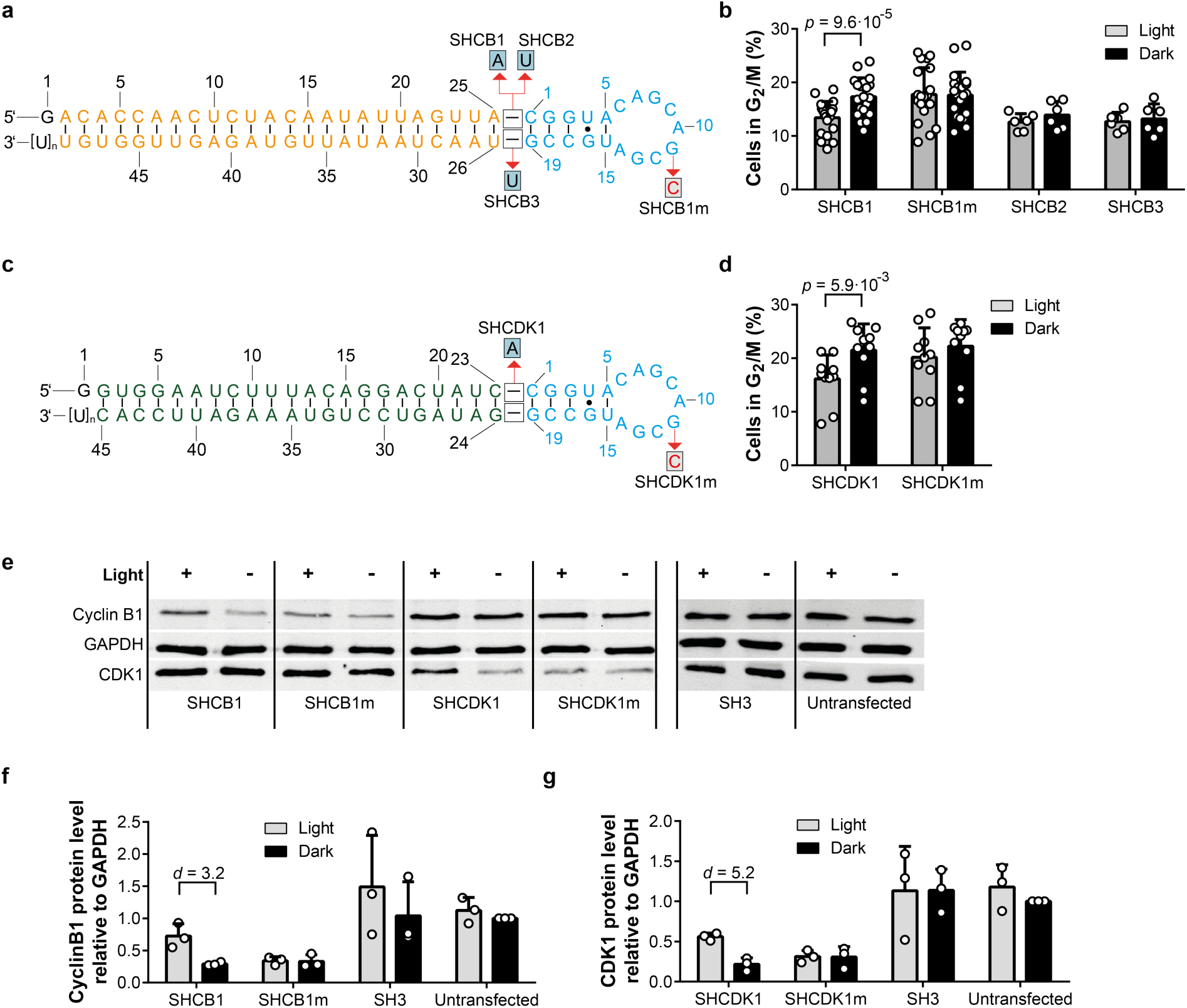
Optoribogenetic control of the mammalian cell cycle. **a**, shRNA variants used to control cyclin B1 gene expression. Blue: aptamer domain; orange: siRNA domain. **b**, Percentages of HEK293PAL cells in G_2_/M phase of the cell cycle when transfected with indicated shRNAs targeting cyclin B1. **c**, shRNA variants used to control CDK1 gene expression. Blue: aptamer domain; green: siRNA domain. **d**, Percentages of HEK293PAL cells in G_2_/M phase of the cell cycle when transfected with indicated shRNAs targeting CDK1. **b,d**, N = at least three biologically independent experiments performed in duplicates. **b**, The identity of SHCB1 and SHCB1m was blinded and double-blinded in one experiment, each. **d**, The identity of SHCDK1 and SHCDK1m was blinded and double-blinded in one experiment, each. **b,d**, Wilcoxon two-sided signed-rank test was used for statistical analysis. **e**, Representative western blot image showing cyclin B1, CDK1 and GAPDH protein expression after transfection with the indicated shRNAs (for complete blots see **Supporting Fig. 21**). **f, g**, Quantification of cyclin B1 and CDK1 protein levels using pixel densitometry (n = three independent experiments). **f, g**, Cohen’s *d* effect size was used for statistical analysis. Values were normalized to non-transfected cells incubated in darkness (Untransfected). **b,d,f,g**, Grey bars: cells incubated under light conditions, black bars: cells incubated under dark conditions. Values are means ± s.d.

## Discussion

In conclusion, we demonstrate the fully genetically encodable light-control of miR and shRNA molecules in mammalian cells. The approach utilizes an aptamer that under blue light binds tightly and specifically to the photoreceptor protein PAL, and this interaction was shown to impact miR and shRNA function in regulating gene expression. We thus created an encoded on-switch, complementing a previously reported off-switch in which the PAL aptamer was embedded directly in the 5’UTR of mRNAs^28^. By offering full genetic encodability, reversibility, and noninvasiveness combined with a small genetic footprint (ca. 1.1 kb), our approach transcends previous approaches for controlling regulatory RNA activity. Specifically, these features distinguish our method from ligand-gated techniques that invariably rely on the exogenous addition of specific compounds, thus abolishing full genetic encoding and limiting their application scope. Our method rivals CRISPR/Cas9-based approaches in its ready adaptability to new target sequences through variation of the modular chimeric RNA. The technology thus unlocks optogenetic control of near-arbitrary gene products at the post-transcriptional level and expands the optogenetic toolbox. Notably, the shRNA-based approach operates dominantly and can hence be used in wild-type cellular backgrounds, thus obviating the laborious construction of transgenic lines. To facilitate adoption of the technology, we investigated in detail sequence determinants affecting the efficiency of light regulation. We demonstrate that single nucleotide variations in the hinge region connecting the miR/siRNA and the aptamer domains impact on regulatory RNA function and allow its fine-tuning, with an up to 15-fold change in protein expression presently. Although we observed a preference for A and U nucleotides of the best-performing shRNAs, we recommend testing all canonical nucleotides (G,U,A, and C) at the hinge region, upstream and downstream of the aptamer domain to identify the most suitable variant. Besides light-dependency, we also demonstrate spatial and temporal regulation and the suitability of the system to control endogenous proteins and cellular behavior, exemplified by controlling the cyclin B1 and CDK1 protein expression. This optoribogenetic approach extends to various shRNA and miR molecules for the investigation of dynamic biological processes by light, e.g., the relationship of proliferation and differentiation of neuronal stem cells, which depends on the progression of the cell cycle^44^. Additionally, optoribogenetic approaches may contribute to the understanding of dynamic micro RNA and protein functions that remain challenging to be resolved with the currently available methodologies.

## Acknowledgement

This work was supported by funds from the European Union ERC (‘OptoRibo’, 615381) to G.M., and the German Research Council (grants MA3442/5-1 and 5-2 to G.M., and MO2192/6-1 to A.M.). We thank Prof. Gruss for critical commenting on the cell cycle data. The FACS core facility of the University Hospital Bonn is acknowledged for support in applying the gating strategy for the eGFP expression experiments.

## Author contributions

S.P. designed the shRNAs, developed and performed all PAL-dependent experiments in mammalian cells and wrote the manuscript. C.M. performed the interaction studies of RNA molecules with PAL, M.C. performed the blinded studies, A.M. conceived the project and discussed experiments, G.M. conceived the study, supervised, discussed, designed the experiments and wrote the manuscript.

## Competing interest statement

The authors declare no competing interests.

## Notes

### Competing Interest Statement

The authors have declared no competing interest.

