## Supporting data for "Optoribogenetic control of regulatory RNA molecules"

**Methods**

**Molecular biology.** All oligonucleotides were purchased from Ella Biotech, Planegg, Germany. Plasmid pIRESneo-FLAG/HA Ago2 was kindly provided by Thomas Tuschl. The plasmids pmCherry-C1, pMetLuc2-Control and peGFP-N1 were purchased from Takara Clontech. All miR and shRNAs described herein were cloned in the pSilencer 2.0-U6 plasmid backbone using restriction cloning. Corresponding miR and shRNA sequences are listed in Supplementary Table S1. The identity of all constructs was confirmed by Sanger DNA sequencing (Eurofins).

**PAL protein purification, *in vitro* transcription and RNA:PAL interaction assay.** For protein expression, the plasmid pET-28c-PAL was transformed into ArcticExpress *e. coli* cells (DE3, Agilent). Bacteria were grown in lysogeny broth (LB) medium supplemented with 50 µg/mL kanamycin, 20 µg/mL gentamycin and 50 µM riboflavin at 37 °C at 120 r.p.m. until an optical density (OD) at 600 nm of 0.6 was reached, at which point expression was induced by addition of 1 mM isopropyl β-D-1-thiogalactopyranoside (IPTG). Incubation continued at 16 °C and 120 r.p.m. for 66 h, after which cells were harvested and lysed by ultrasound. The lysate was cleared by centrifugation and applied to an immobilized nickel ion affinity column (Marcherey Nagel). Protein was eluted in buffer A (50 mM Tris/HCl, 200 mM NaCl, 200 mM imidazole, 10 % w/v glycerol, pH 7.6). Protein was dialyzed into buffer B (12 mM HEPES/KOH, pH 7.2, 135 mM KCl, 10 mM NaCl, 10 % w/v glycerol). Identity of the protein was verified by polyacrylamide gel electrophoresis (PAGE). Protein concentration was determined by absorption spectroscopy using an extinction coefficient of 12,500 M^-1^ cm^-1^ at 447 nm.

*In vitro* transcription was performed using T7 RNA polymerase. DsDNA template coding for shRNA sequences was modified upstream by implementing T7 promoter sequence and two additional guanine residues (bold) after the transcription start site to ensure transcription (TAATACGACTCACTATAG**GG**, underlined G: transcription start site). DNaseI (Roche) digested RNA was purified by PAGE and recovered by electroelution (8 M ammonium acetate, 45 min, 150 V). PAGE and electroelution were performed using TBE buffer (100 mM Tris, 100 mM boric acid, 2 mM ethylenediaminetetraacetic acid, pH 8.0).

For RNA:PAL interaction assays, PAL protein was biotinylated with a four-fold excess of EZ-Link Sulfo-NHS-LC-Biotin according to the manufacturer’s instructions (Thermo Fisher Scientific) and coupled to streptavidin-coated wells of a plate (Pierce Streptavidin Coated Plates, Black, 96-Well, Thermo Fisher Scientific). Wells were washed three times with 200 µl buffer B. Subsequently, 100 µl of 1.5 µM biotinylated PAL in buffer B was added and the coupling was performed in darkness over night at 4 °C. Afterwards, wells were washed three times with 200 µl buffer B. *In vitro* transcribed *pre*-miR or shRNA constructs were incubated with immobilized PAL at the indicated concentrations in 100 µl buffer B for 30 min at 25 °C under light (λ_max_= 465 nm, 2.15 W/cm^2^) or in darkness, followed by three washing steps with 200 µl buffer b for 3 min each. Fluorescence detection was performed by adding 150 µl RiboGreen reagent (Quant-iT RiboGreen RNA Reagent, Thermo Fisher Scientific), diluted 500-fold in 1x TE buffer (10 mM Tris/HCl, pH 7.5, 1 mM ethylenediaminetetraacetic acid) to each well. After 1 h incubation in darkness, fluorescence intensity was measured on a Tecan Ultra plate reader (Tecan) at excitation and emission wavelengths of 500 and 525 nm, respectively. Results were normalized to full length aptamer (53) incubated under light conditions.

**LED array.** Blue-light was administered to cells in pulses (30 sec light on, 30 sec light off) by a custom LED array (illuminance at 106 µW/cm^2^) with λ_max_= 465 nm. The LED array was powered using a custom built microcontroller. Cells were exposed to light immediately after transfection until they were subjected to further investigation.

**Light plate apparatus (LPA).** A custom-made replicate of the LPA was produced by Hanns-Martin-Schmidt and used for the photomask experiment.

**Working with mammalian cell lines.** HEK293 cells (CLS Cell Lines Service) were cultured in DMEM medium (high glucose, GlutaMAX), supplemented with 1 % non-essential amino acids (NEAA), 1 % Sodium Pyruvate (Thermo Fisher Scientific) and 10 % fetal calf serum (FCS, Sigma-Aldrich), at 37 °C in a humidified 5 % CO_2_ atmosphere and were passaged every 2–3 days. HEK293PAL cells were cultured one week before and after cell sorting with 1 % penicillin/streptomycin and in presence of Geneticin (G418, 400 µg/mL, Thermo Fisher Scientific). Mycoplasma testing was performed every three months using PCR detection (Minerva biolabs).

**Transient transfection.** Cells were transfected 24 h after incubation in darkness in 500 µl DMEM supplemented with 1 % NEAA and 1 % Sodium Pyruvate. 1.5 µL Lipofectamine2000 (Thermo Fisher Scientific) was used per well in a 24-well plate format. For transfection experiments involving AGO2, cells were transfected with 250 ng plasmid DNA (e.g., pSilencer, AGO2 and pEGFP-N1 plasmid at a mass ratio of 2:2:1). For reporter assays without AGO2 overexpression, cells were transfected with 500 ng plasmid DNA (e.g. pSilencer and pEGFP-N1 or PMetLuc2-Control plasmid at a mass ratio of 100:1). For transfections of shRNAs targeting intrinsic mRNAs, 500 ng of the corresponding pSilencer plasmid variant was transfected. Plasmids and Lipofectamine2000 were each diluted in 50 µl Opti-MEM (Thermo Fisher Scientific) per well and incubated 5 min before mixing at room temperature. After 20 min further incubation at room temperature, 100 µl transfection mix was added per well. Four hours after transfection, 60 µl FCS were added per well.

**Flow cytometry analysis.** Flow cytometry was performed on a BD FACSCanto II (BD Biosciences). Data was processed using FlowJo version 9.6.3 software. Cells were initially gated with SSC and FSC channels for cell debris and single cell populations. For measuring eGFP, cells were additionally gated using the fluorescein isothiocyanate (FITC) channel to determine eGFP positive cells. For cell cycle measurements, cells were additionally gated using Phycoerythrin (PE) channel to determine the percentage of cells in the different cell cycle phases. At least 30.000 cells were analyzed from each sample. eGFP and propidium iodide were excited with a 488 nm laser and detected with a 530/30 or 585/42 filter set, respectively. mCherry was excited with a 633 nm laser and detected with a 660/20 filter set.

**Isolation of monoclonal cells.** For the generation of stable monoclonal HEK293 cell lines expressing mCherry-PAL (HEK293PAL), 10^6^ cells were seeded into each well of a 6 well plate and transfected by Lipofectamine2000 transfection on the following day using 2.5 µg plasmid DNA and 8 µl Lipofectamine2000. Four hours after transfection, the supernatant was discarded and cells were washed with PBS before the addition of 3 mL cell medium. After three days, the medium was supplied with 400 µg/mL G418. One week before and after cell sorting cell medium was supplemented with 1 % penicillin/streptomycin (Thermo Fisher Scientific). After five weeks of selection, single cells, which strongly express mCherry, were sorted *via* fluorescence activated cell sorting into a well of a 96-well plate. Cells were cultivated until 80 % confluency was reached in a T-175 culture flask. Then, cells were frozen or used in further experiments. We did not notice any change in growth rate and cell fitness of the HEK293PAL cell line compared to HEK293 cells.

**Luciferase reporter assays in mammalian cells.** 1 × 10^5^ HEK293PAL cells were seeded in two separate 24-well plates per well. After 24 h, cells were transfected according to the protocol described above. Transfected cells were incubated for 19 h in the presence of blue light using the LED array or in darkness.

For reversibility assays, one control plate was kept constantly in darkness. Another plate was incubated under varying light conditions after transfection as indicated by the time lines shown in **Fig. 1h** and **k**. Immediately before the light irradiation status was altered for one plate, cell medium was exchanged for both plates involved in the assay (plate incubated constantly in darkness and plate exposed to varying light conditions).

For the luciferase assay, 50 μl of the cell culture supernatant was transferred to wells of a white 96-well plate (LUMITRAC 200, Greiner). 5 μl luciferase substrate dissolved in buffer according to the manufacturer’s instructions (Ready-To-Glow Secreted Luciferase, Takara Clontech) was added and the reaction was incubated for 3 min at room temperature. The luminescence signal was measured using an EnSpire plate reader (PerkinElmer) with an integration time of 5 s.

For the static luciferase assay (**Fig. 1b**), values were normalized to control *pre*-miR transfection (SHD) incubated in darkness, where no influence on luciferase expression was expected. For the reversibility luciferase assay (**Fig. 1i**, **l**), values were normalized to aptamer point mutant *pre*-miR21 variant (SHC) incubated constantly in darkness, where no light-dependency was expected. Normalization was performed to each time point.

**Photomask experiment.** 7.5 × 10^4^ HEK293PAL cells were seeded in black 24-well plates with clear bottom (VisionPlate, 4titude). After 24 h, cells were transfected using the protocol for AGO2 overexpression as described in the transient transfection section. Then, the plate was mounted onto the custom-made LPA device and irradiated with 10 µW/cm^2^ of constant light (λ_max_= 465 nm). After 48 h, cells were analyzed by confocal laser scanning microscopy (LSM 710) using a 10x/0.45 objective and image concatenation (10% overlay). Imaging was performed at 37 °C. eGFP fluorescence was visualized as green color and image histograms were adjusted to 5/10 before .tiff picture export (Zen Black software, Zeiss). Image brightness was adjusted to + 150 and image sizes were adjusted to 300 x 300 pixels using Adobe Photoshop CS5 software.

**eGFP reporter assays in mammalian cells**. 1 × 10^5^ HEK293PAL cells were seeded in two separate 24-well plates per well. After 24 h, transfection was performed as indicated in the transient transfection section. For the miR21-based reporter assays, the 3’UTR of peGFP-N1 bearing miR21 binding sites (**Supporting Figure 1b**) was used. Cells were incubated for 44 h in the presence of blue light using the LED array or in darkness. Then, the cell supernatant was aspirated, cells were washed and resuspended in PBS (25 °C). The subsequent analysis was performed using flow cytometry. The percentage of eGFP positive cells was normalized to control *pre*-miR transfection (SHD) incubated in darkness where no influence on eGFP expression was expected for miR experiments. For shRNA experiments, the percentage of eGFP positive cells was normalized to functional *pre*-miR21 transfection containing the PAL aptamer (SHA) and incubated in darkness as no influence on eGFP expression was expected due to the absence of miR21 binding sites in the 3’UTR sequence of the eGFP mRNA.

**Fluorescence microscopy of HEK293PAL cells.** 5 × 10^4^ cells were seeded in black 24-well plates with clear bottom (μ-plate, ibidi). After 24 h, cells were transfected using the protocol for AGO2 overexpression as described in the transient transfection section. Cells were incubated for 44 h in the presence of blue light (465 nm, 106 μW cm^−2^, 30 s pulses) or in darkness. Then, the supernatant was replaced by cell medium containing 5 µg/mL Hoechst 33342. After 10 min incubation at 37 °C, the supernatant was replaced with cell medium only. Next, cells were analyzed by confocal laser scanning microscopy (LSM 710, Zen Black software, Zeiss) using a 20x/0.8 objective. We generated pictures comprising of 1528 x 1528 pixels with a resolution of 0.19 µm per pixel and a pixel dwell time of 2.11 µs for imaging of each fluorophore. Fluorescence of mCherry (excitation (ex)/emission (em): 543/578–696 nm), Hoechst 33342 (ex/em: 405/410-494) and eGFP (ex/em: 488/494-574 nm) was monitored, respectively. Imaging was performed at 37 °C.

**Western blot analysis.** HEK293PAL cells were lysed 44 h after transfection in RIPA buffer (Thermo Fisher Scientific) containing 1 mM Phenylmethylsulfonyl fluoride. Lysates were cleared by centrifugation (14,000 g for 15 min, 4°C). Cleared lysates in Laemmli buffer were incubated at 95 °C for 5 min before loading on SDS-PAGE gels. Protein quantification was performed using the Pierce BCA protein assay kit according to the manufacturer’s instruction (Thermo Fisher Scientific). 5 µg of protein per lane was loaded onto 10 or 12,5 % SDS-PAGE gels and blotted in Transfer Buffer (2.5 mM Tris, 2 % (w/v) glycine, 0.9 M urea) onto a nitrocellulose membrane (GE Healthcare Life Sciences) using a Bio-Rad Trans-Blot SD Semi-Dry Transfer Cell (BioRad) for 75 min at 20 V and 30 W. Membranes were blocked with TBS-T buffer (20 mM Tris/HCl, pH 7.6, 150 mM NaCl, 0.05 % Tween 20 (v/v)) containing 5 % BSA (AppliChem, Western Blot grade) under agitation at room temperature for 1 h. Blots were cut according to the protein ladder (Prestained Protein Ladder – Mid-range molecular weight (10 – 180 kDa), abcam) in a way that all target proteins can be individually incubated with the respective primary antibody (mouse *anti*-cdc2 (CDK1), Cell signaling POH1, #9116 (1:1000); mouse *anti*-GAPDH, Santa Cruz Biotechnology sc-47724 (1:4000); goat *anti-*Cyclin B1, R&D Systems AF6000 (1:1000)) at 4 °C overnight or at room temperature for 1 h in TBS-T containing 5 % BSA. Detection was performed with IRDye 800CW goat *anti*-mouse (Li-cor 926-32210), donkey *anti*-goat (Li-cor 926-32214) and goat *anti*-rabbit (Li-cor 926-32211) at a dilution of 1:15000, respectively.

Antibody-stained blot pieces were arranged, and fluorescence images were acquired using an Odyssey Imaging System’s (Li-cor) 800 nm channel (ex/em: 785/810 nm) to visualize bound 800CW secondary antibodies. Pixel densitometry of blot bands was performed using Fiji software (ImageJ) by creating rectangles of equal sizes for each sample lane followed by quantification of the area under the peak of each protein spot. Relative density was calculated by dividing values obtained from putative CDK1 or cyclin B1 protein bands through the values obtained from the putative GAPDH band of each lane. Relative density values were normalized to untransfected cells incubated in darkness.

**Cell cycle assay.** 1 × 10^5^ HEK293PAL cells per well were seeded in two separate 24-well plates. 24 h after seeding, transfection was performed according to the transfection protocol for shRNAs targeting intrinsic mRNAs as indicated in the transient transfection section. Cells were incubated for 44 h in the presence of blue light (465 nm, 106 μW cm^−2^, 30 s pulses) or in darkness. 44 h after transfection, cells were fixed in 70% ice-cold Methanol in PBS and incubated for at least 30 min at 4 °C followed by RNAse A (50 µg/mL) and propidium iodide (PI, 50 µg/mL) treatment for 30 min at 37 °C under mild agitation. Subsequent analysis of cell cycle distribution quantified by PI fluorescence per cell was performed using flow cytometry.

**mCherry quantification in HEK293PAL cells.** 1 × 10^6^ HEK293PAL cells were lysed in 250 µl RIPA buffer (Thermo Fisher Scientific) containing 1 mM Phenylmethylsulfonyl fluoride (PMSF). Lysates were cleared by centrifugation (14,000 g for 15 min, 4°C). Generation of a mCherry standard curve and fluorescence measurements were performed using mCherry Quantification Kit (BioCat) according to the manufacturer’s instruction with the exception that RIPA buffer containing 1 mM PMSF was used instead of mCherry Assay Buffer. For calculation of the concentration of mCherry-PAL a cellular volume of 4000 µm^3^ was assumed and the cytoplasm covering 1/3 of this volume (Mateus *et al.*, *Molecular Pharmaceutics*, 2013).

**Determination of 3’ ends of artificial miR21-5p.** 1 × 10^5^ HEK293PAL cells were seeded in a 24-well plate. 24 h later, transfection with SHA and luciferase reporter plasmid was performed and cells were further incubated in darkness. 19 h after transfection total RNA extraction with TRIzol was performed according to the manufacturer’s instructions (Thermo Fisher Scientific). 1,5 µg total DNase-treated RNA was poly-adenylated using Poly-A Polymerase according to the manufacturer’s instructions (New England Biolabs). The reaction was purified by phenol/chloroform extraction followed by ethanol precipitation. Reverse transcription was performed using 1 µg poly-adenylated cDNA and Bioscript Reverse Transcriptase (RT-Primer: ATTCTAGAGGCCGAGGCGGCCGACATGTTTTTTTTTTTTTTTT-TTTTTTTTTTTTTT) according to the manufacturer’s instructions (Bioline). After another round of phenol/chloroform extraction followed by ethanol precipitation, PCR amplification was performed using Taq Polymerase (Forward Primer: CGCCtagcttatcagactgatgt, reverse Primer: ATTCTAGAGGCCGAGGCGGCCGACATG). Cloning was performed using TOPO-TA cloning kit according to the manufacturer’s instructions (Thermo Fisher Scientific). Plasmids were isolated from individual clones using Plasmid DNA purification kit (Marcherey-Nagel) and sent for sequencing (Eurofins).

**Calculation of fold changes.** Light-dependent fold changes were calculated by dividing values (crude values for luciferase assay, percentage of eGFP positive cells for eGFP assays) of samples incubated under light conditions through samples incubated in darkness (duplicates) to obtain four values for each independent experiment.

**Blinded experiments.** The identity of SHCB1, SHCB1m, SHCDK1, SHCDK1m and SH3 which are shown in **Fig. 3b**, **d** and **Supporting Figure 7** and **9** were blinded and double-blinded in one experiment, each. For the blinded experiment, the identity of these PSilencer plasmids have been blinded before transfection by a second person. For the double-blinded experiment, another experimenter performed the assay that was blinded by a third person in a similar way. Identity of the samples was confirmed after data evaluation between the experimenter and the person who performed the blinding.

**Statistics and reproducibility.** Prism 6.01 (GraphPad Software, Inc.) was used to generate graphs and calculate *P* values. For all statistical analysis, no gaussian distributions were assumed due to limited sample sizes. Wilcoxon two-sided signed-rank test was used to compare equally treated cell samples incubated under the indicated light conditions. Therefore, a paired observation was assumed. Two-sided Mann-Whitney *U* test was used to compare light-dependent fold changes between differently treated groups (e.g. different time points). Therefore, an unpaired observation was assumed. Cohen’s *d* effect size was used to calculate effect sizes in Western Blot analysis derived from equally treated cell samples incubated under the indicated light conditions. In this case, sample size was too small to test for significance. Datasets are presented as mean ± s.d., if not otherwise stated.

**Reporting Summary.** Further information on research design is available in the Nature Research Reporting Summary linked to this article.

**Data availability.** Additional raw data or materials are available from the corresponding authors upon request.

**Biological material.** HEK293PAL cell line and plasmids is available from the corresponding authors upon request.

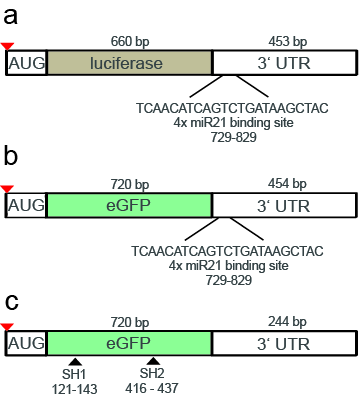

**Supporting Figure 1: Schematics of regulatory RNA binding sites on luciferase and eGFP reporter mRNA molecules.** Schematic representations of luciferase (**a**) and eGFP mRNA (**b**) coding regions (brown (**a**), green(**b**)) and corresponding 3’ untranslated regions (3’UTRs) with four miR21 binding sites were used as reporter mRNAs for *pre*-miR21 experiments (**Fig. 1**). **c**, Schematic representation of eGFP reporter mRNA for shRNA experiments (**Fig. 2**) with no miR21 binding sites embedded in the 3’UTR. **c**, Putative binding sites of siRNAs originating from SH1 and SH2 are indicated by black arrows. **a**-**c**, Numbers indicate base numbers relative to translation to start codon as well as the length of protein coding regions and the 3’UTR regions.

5’GCGGCCGCTCAACATCAGTCTGATAAGCTACTAATCAACATCAGTCTGATAAGCTACTAATCAACATCAGTCTGATAAGCTACTAATCAACATCAGTCTGATAAGCTAGCGGCCGCgactctagatcataatcagccataccacatttgtagaggttttacttgctttaaaaaacctcccacacctccccctgaacctgaaacataaaatgaatgcAATTGACAGCCCATCGACTGGTGTTGCTAAACAGCCCATCGACTGGTGTTGCTAAACAGCCCATCGACTGGTGTTGCTAAACAGCCCATCGACTGGTGTTGcAATTG3’

**Supporting Figure 2: DNA sequence of the miR21 binding sequences located in the 3’UTR of the reporter plasmids.** Four binding sites (blue) complementary to miR21-5p were cloned into the NotI restriction site (grey). Four binding sites (green) complementary to miR21-3p were cloned into the MfeI restriction site (grey).

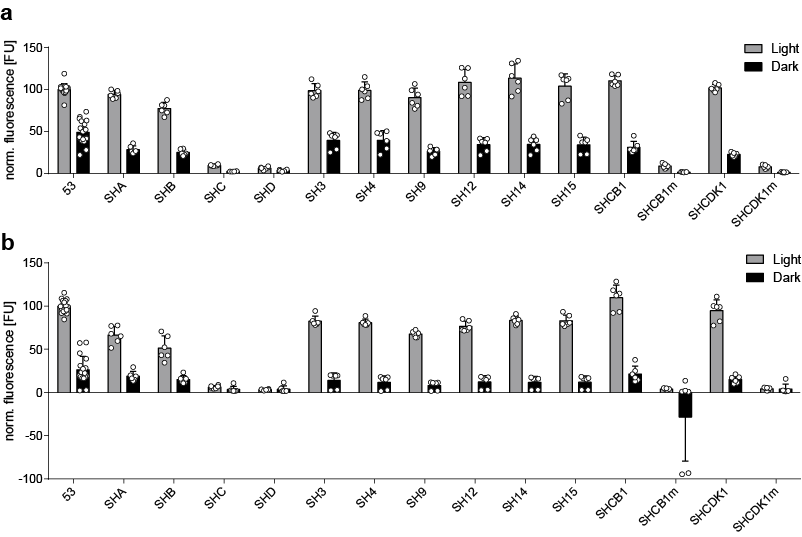

**Supporting Figure 3: Aptamer-conjoined regulatory RNAs bind to PAL light-dependently *in vitro*, whereas binding is reduced for respective point mutants.** Biotinylated PAL protein was immobilized on streptavidin coated wells. Binding of 1000 nM (**a**) or 100 nM (**b**) *pre*-miR21, eGFP shRNA or cell cycle regulatory RNA constructs was quantified in presence of 0.5 mg mL^-1^ heparin and 0.5 mg mL^-1^ BSA by RiboGreen fluorescence. **a**,**b**, Values are processed by subtracting background fluorescence from equally treated wells without immobilized PAL and subsequent normalization to 53 incubated under light conditions. **a**,**b**, N = three independent experiments performed in duplicates. Grey bars: light conditions, black bars: darkness. Values are means ± s. d.

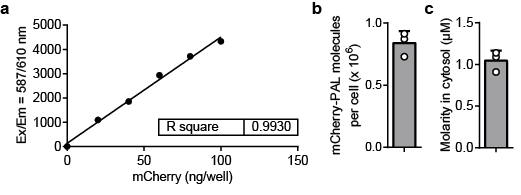

**Supporting Figure 4: Quantification of mCherry-PAL expression levels in the transgenic cell line HEK293PAL.** **a**, Representative standard curve for determination of the linear range of mCherry quantification. **b**, Determination of average mCherry-PAL molecules per cell using the corresponding mCherry standard curve (n = three biological replicates). **c**, mCherry-PAL molarity in the cytosol calculated from (**b)**. **b**,**c**, N = three independent experiments performed in duplicates. Values are means ± s. d.

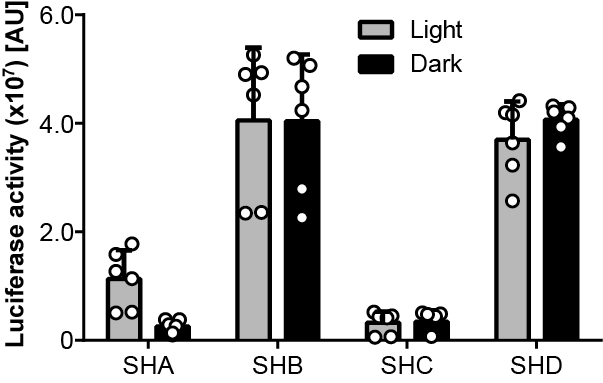

**Supporting Figure 5: A *pre*-miR21-aptamer chimera enables light-control of luciferase activity.** Luciferase activity after transient transfection of the indicated *pre*-miR21 variants. HEK293PAL cells were incubated under the indicated light conditions prior to measurement. A normalized variant of this dataset is shown in **Fig. 1b.** Experiment was performed in duplicates and three independent replicates. Grey bars: light conditions, black bars: dark conditions. Values are means ± s.d.

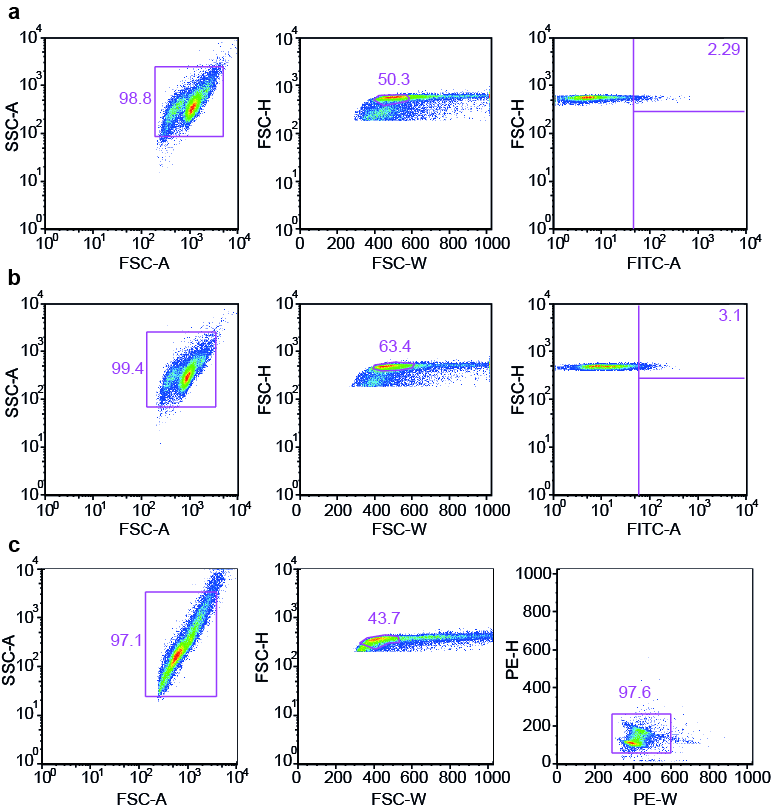

**Supporting Figure 6: Gating strategies to identify eGFP positive cells for miR21 (a), shRNA experiments (b) and cell cycle phases (c). a-c**, Cell debris was excluded using side scatter area (SSC-A) *vs*. forward scatter area (FSC-A). Singlet cells were detected using forward scatter height (FSC-H) *vs*. forward scatter width (FSC-W). For miR21 and shRNA experiments (**a**, **b**), eGFP positive cells were identified using FSC-H *vs*. Fluorescein isothiocyanate area (FITC-A). **a**, Cells transfected with SHA and incubated in darkness were set to 2.29 % eGFP positive cells and gating was applied to all other tested samples. **b**, Cells transfected with SH1 and incubated in darkness were set to 3.1 % eGFP positive cells and gating was applied to all other tested samples. **c**, Cell cycle debris and apoptotic cells were excluded using Phycoerythrin height (PE-H) *vs*. Phycoerythrin width (PE-W). Flow cytometry was performed using a BD FACS Canto II instrument (BD Bioscience). Data processing was performed using FlowJo (9.6.3) and GraphPad Prism (6.01).

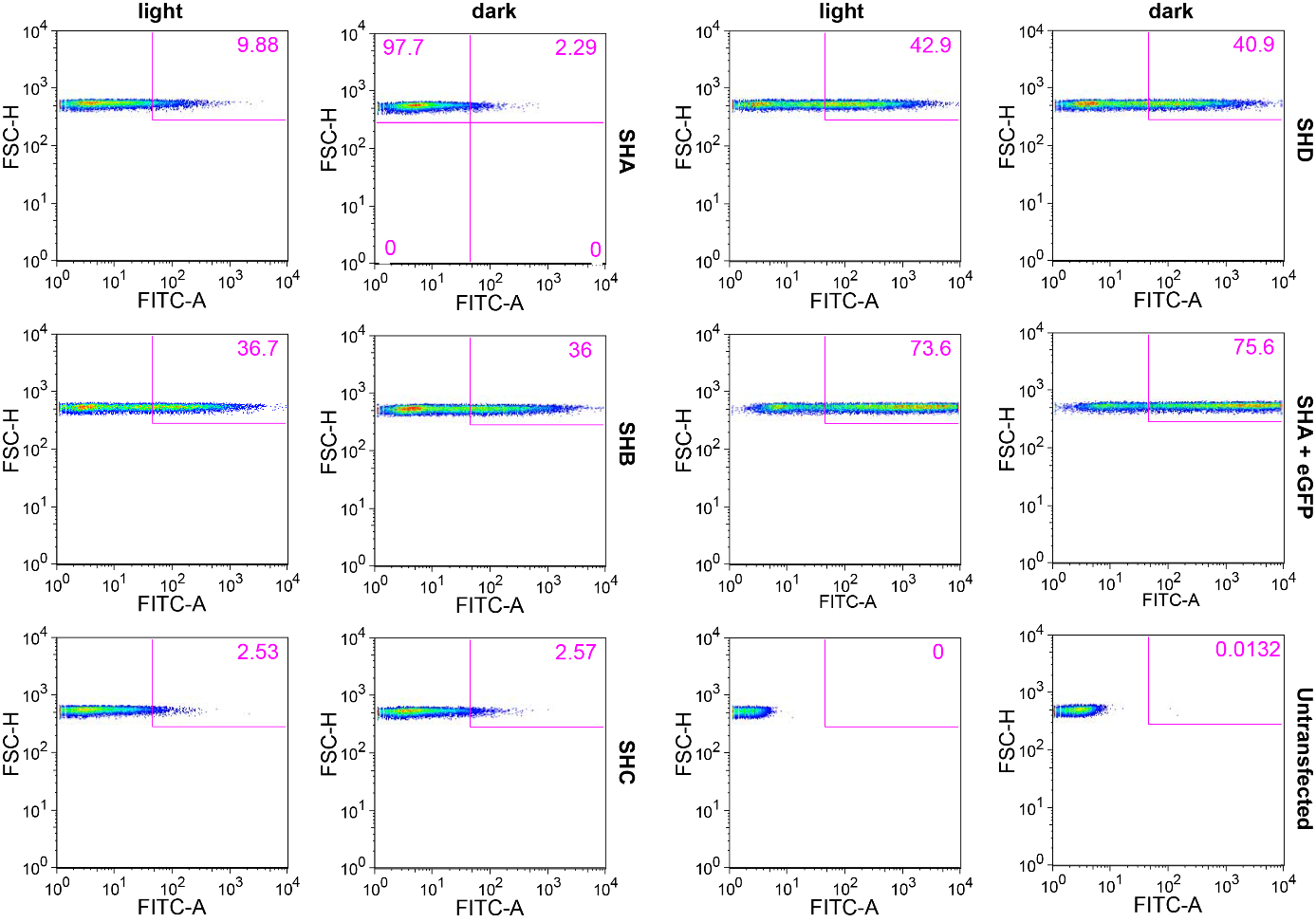

**Supporting Figure 7: Detection of eGFP expression for miR21 experiments in absence of AGO2 overexpression shown in Fig. 1d.** Representative flow cytograms of HEK293PAL cells after transfection of the indicated *pre*-miR variants. Gating strategy was applied as outlined in **Supporting Figure 6a**. HEK293PAL cells were incubated under the indicated light conditions prior to measurement. eGFP expression was detected by using forward scatter height (FSC-H) *vs*. Fluorescein isothiocyanate area (FITC-A) channel. Flow cytometry was performed using a BD FACS Canto II instrument (BD Bioscience). Data processing was performed using FlowJo (9.6.3) and GraphPad Prism (6.01).

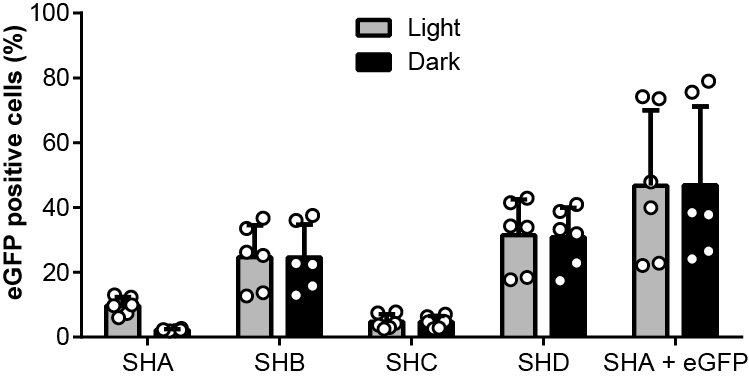

**Supporting Figure 8: A *pre*-miR21-aptamer chimera enables light-control of eGFP expression.** Number of eGFP positive cells after transient transfection of the indicated *pre*-miR21 variants. HEK293PAL cells were incubated under the indicated light conditions prior to measurement. A normalized variant of this dataset is shown in **Fig. 1d.** eGFP expression was identified as indicated in **Supporting Fig. 6a**. Experiment was performed in duplicates and three independent replicates. Grey bars: light conditions, black bars: dark conditions. Values are means ± s.d.

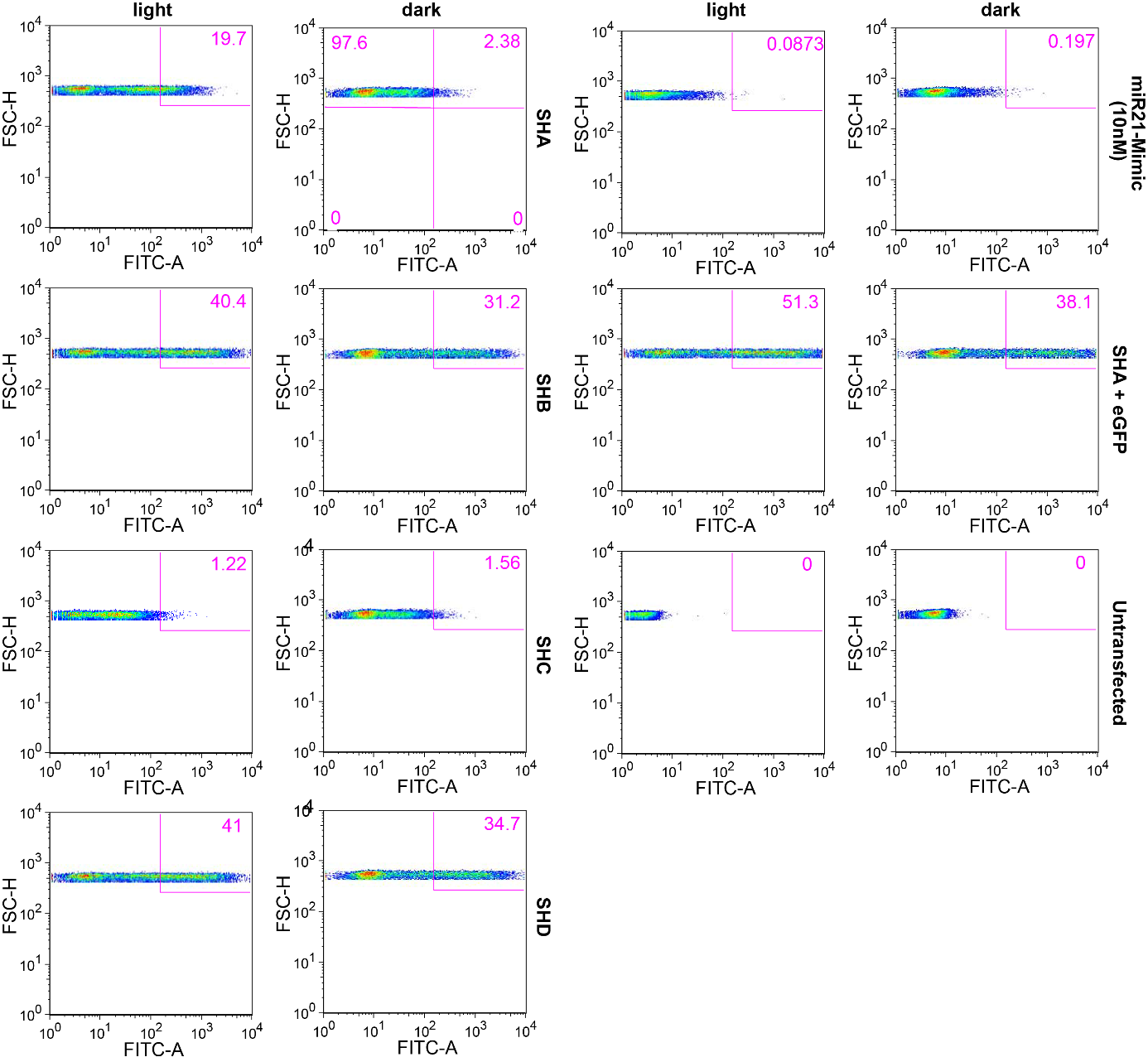

**Supporting Figure 9: Detection of eGFP expression for miR21 experiments in presence of elevated levels of AGO2 shown in Fig. 1f.** Representative flow cytograms of HEK293PAL cells after transfection of the indicated *pre*-miR variants. Gating strategy was applied as outlined in **Supporting Figure 6a**. HEK293PAL cells were incubated under the indicated light conditions prior to measurement. eGFP expression was detected by using forward scatter height (FSC-H) *vs*. Fluorescein isothiocyanate area (FITC-A) channel. Flow cytometry was performed using a BD FACS Canto II instrument (BD Bioscience). Data processing was performed using FlowJo (9.6.3) and GraphPad Prism (6.01).

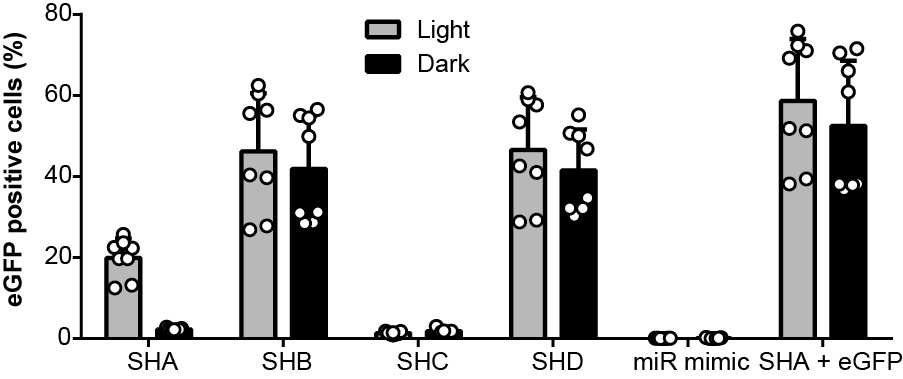

**Supporting Figure 10: Elevated levels of AGO2 increase light-control of eGFP expression mediated by a *pre*-miR21-aptamer chimera.** Number of eGFP positive cells after transfection of the indicated *pre*-miR21 variants or 10 nM miR21 mimic. A normalized variant of this dataset is shown in **Fig. 1f.** eGFP expression was identified as indicated in **Supporting Fig. 6a**. Experiment was performed in duplicates and four independent replicates. Grey bars: light conditions, black bars: dark conditions. Values are means ± s.d.

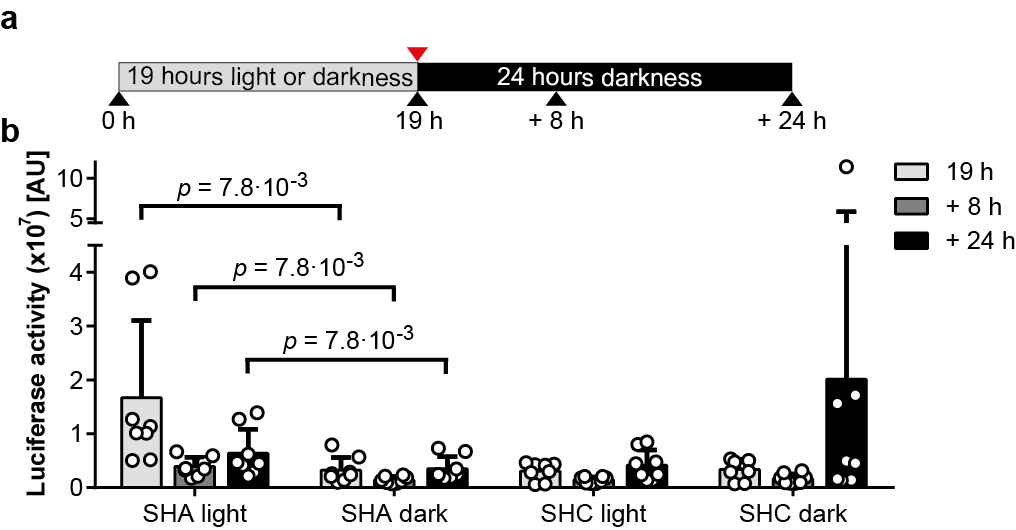

**Supporting Figure 11: Luciferase activity is reversible. a**, Illumination protocol applied in (**b**). Red arrow: time point of medium exchange, black arrows, time points of sample collection. **b**, Luciferase activity after transient transfection of the indicated *pre*-miR21 variants. A normalized variant of this dataset is shown in **Fig. 1i**, in which values were normalized to aptamer point mutant *pre*-miR21 variant (SHC) incubated in darkness, where no light-dependency was expected. Normalization was performed to each time point. HEK293PAL cells were either incubated under conditions shown in (**a**) or constantly in darkness prior to measurement. **b**, Wilcoxon two-sided signed-rank test was used for statistical analysis as a paired observation was assumed. Experiment was performed in duplicates and four independent replicates. Values are means ± s.d.

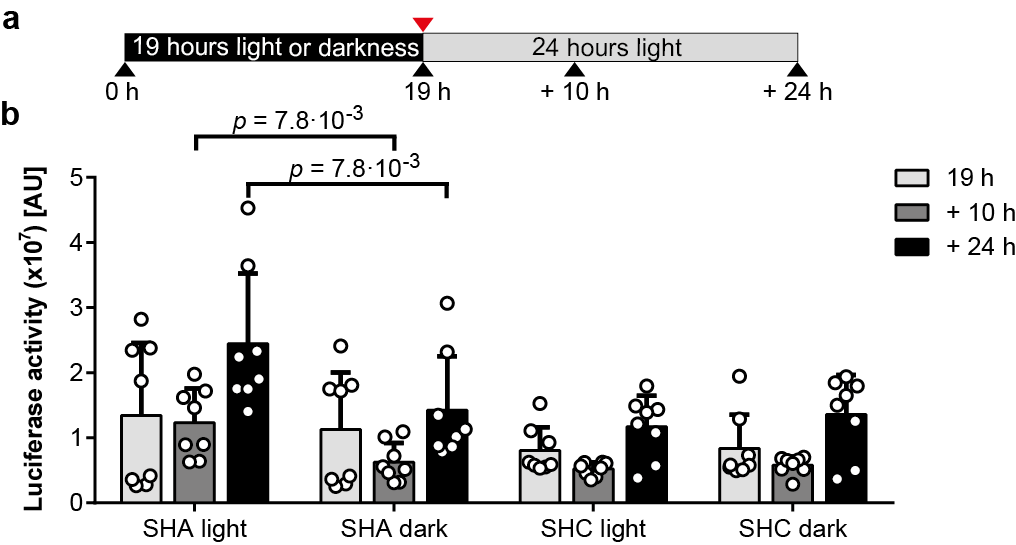

**Supporting Figure 12: Luciferase activity is inducible. a**, Illumination protocol applied in (**b**). Red arrow: time point of medium exchange, black arrows, time points of sample collection. **b**, Luciferase activity after transient transfection of the indicated *pre*-miR21 variants. A normalized variant of this dataset is shown in **Fig. 1l**, in which values were normalized to aptamer point mutant *pre*-miR21 variant (SHC) incubated in darkness, where no light-dependency was expected. Normalization was performed to each time point. HEK293PAL cells were either incubated under conditions shown in (**a**) or constantly in darkness prior to measurement. **b**, Wilcoxon two-sided signed-rank test was used for statistical analysis as a paired observation was assumed. Experiment was performed in duplicates and four independent replicates. Values are means ± s.d.

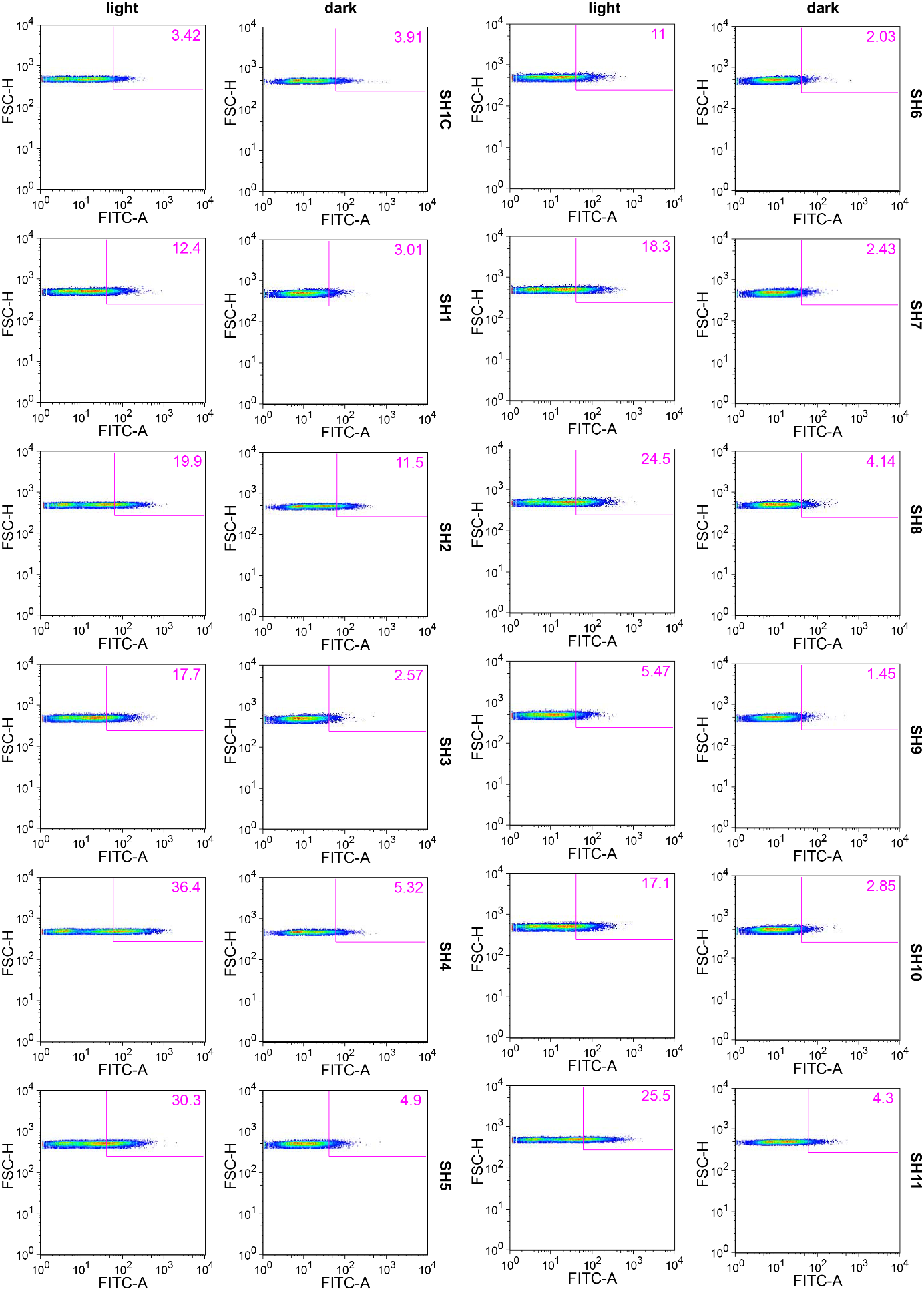

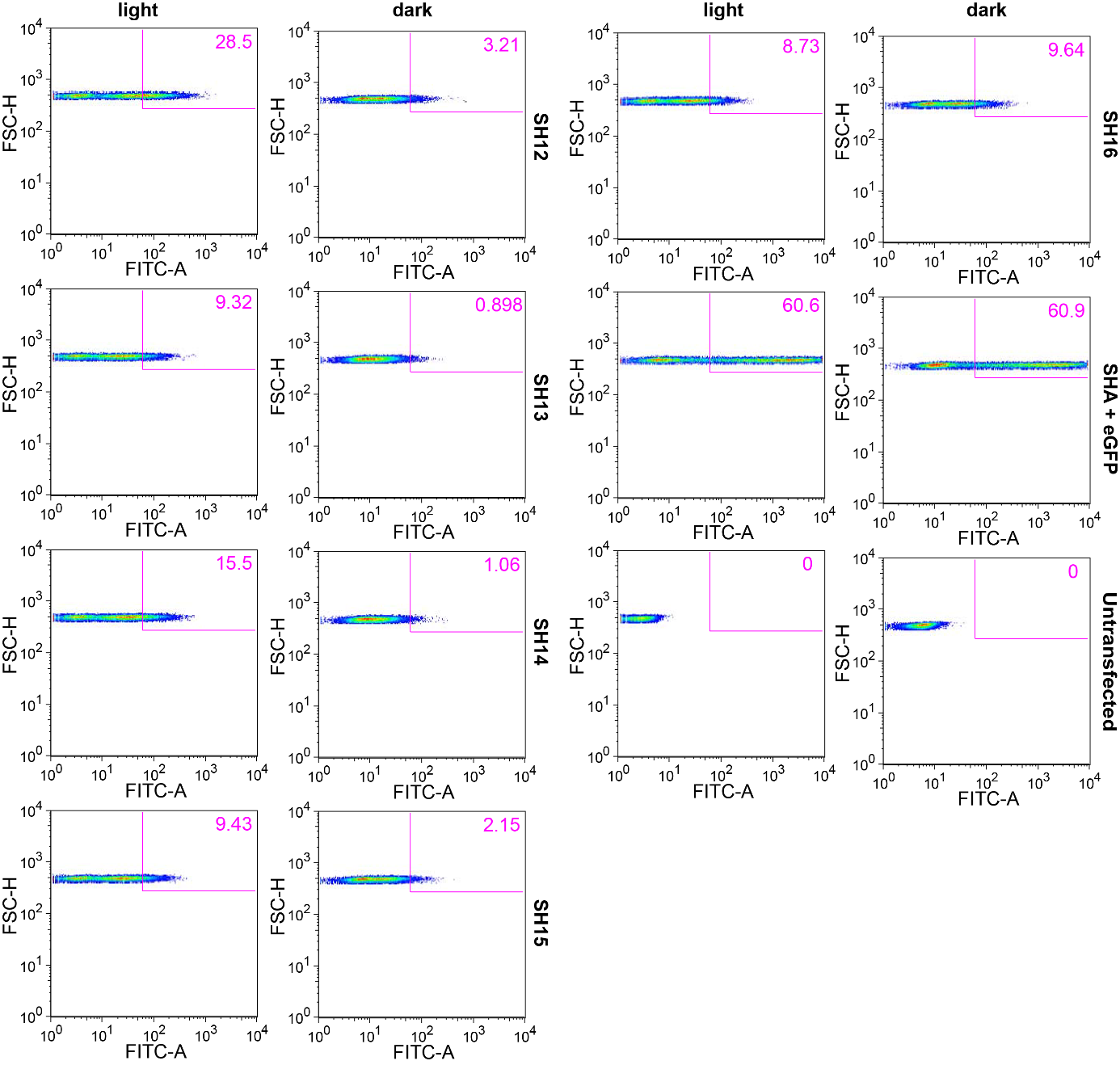
**Supporting Figure 13: Detection of eGFP expression for shRNA experiments.** Representative flow cytograms of HEK293PAL cells after transfection of the indicated shRNA variants. Experiment is shown in **Fig. 2c**,**d**,**f**,**g**,**i**,**j** of the main text. Gating strategy was applied as outlined in **Supporting Figure 6b.** HEK293PAL cells were incubated under the indicated light conditions prior to measurement. eGFP expression was detected by using forward scatter height (FSC-H) *vs*. Fluorescein isothiocyanate area (FITC-A) channel. Flow cytometry was performed using a BD FACS Canto II instrument (BD Bioscience). Data processing was performed using FlowJo (9.6.3) and GraphPad Prism (6.01).

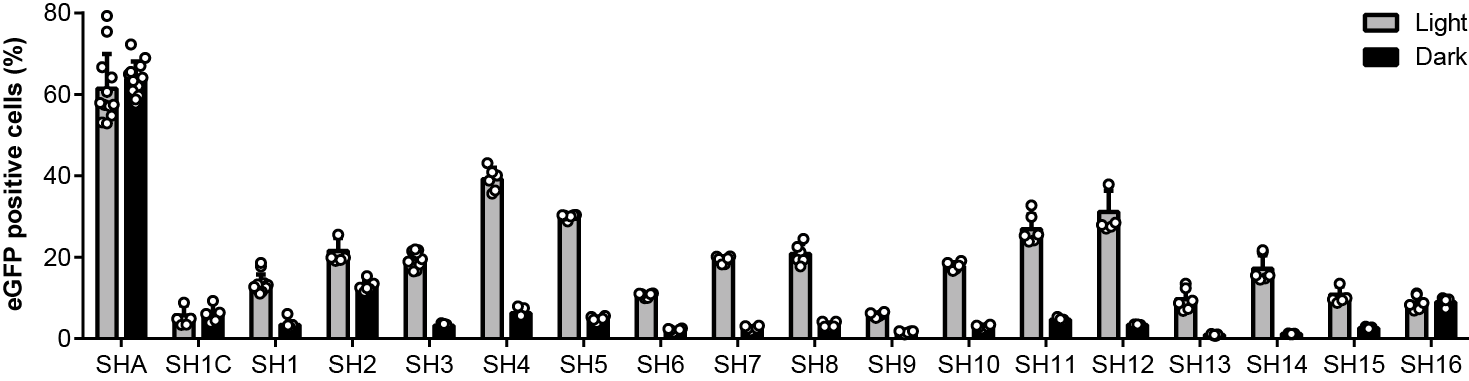

**Supporting Figure 14: shRNA-aptamer chimeras enable light-control of eGFP expression.** Number of eGFP positive cells after transient transfection of AGO2 and the indicated shRNA variants. HEK293PAL cells were incubated under the indicated light conditions prior to measurement. eGFP expression was identified as indicated in **Supporting Fig. 6a**. A normalized variant of this dataset is shown in **Fig. 2c**,**f**,**i.** Experiment was performed in duplicates and at least three independent replicates. Grey bars: light conditions, black bars: dark conditions. Values are means ± s.d.

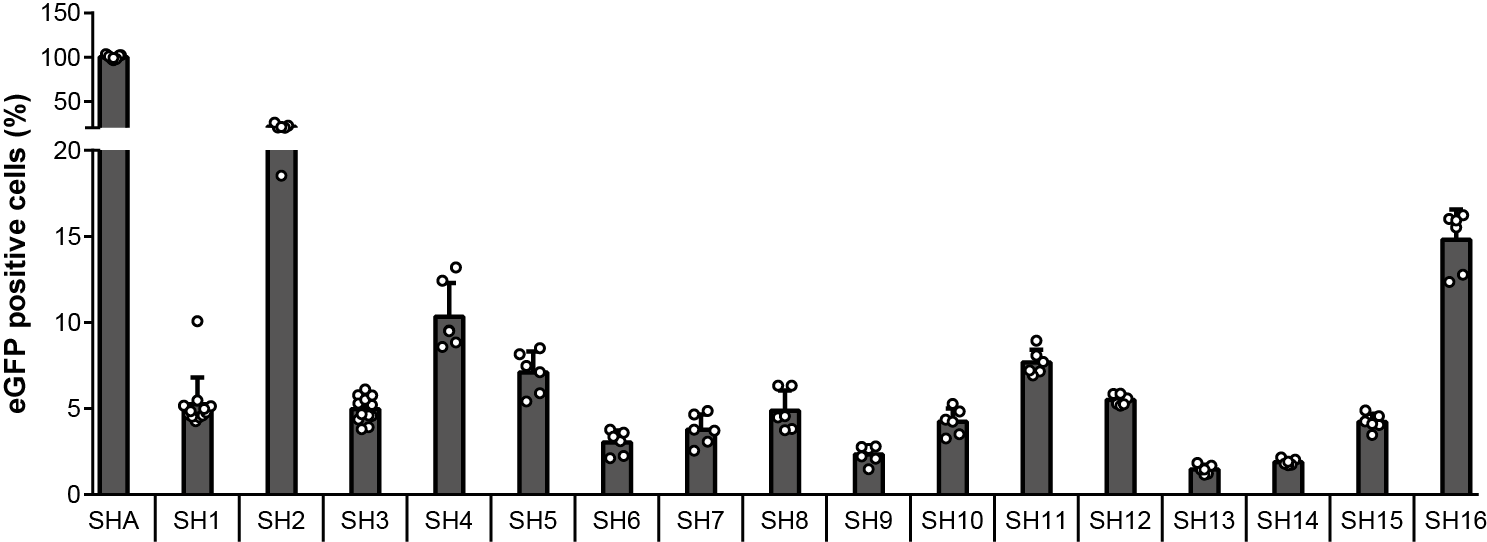

**Supporting Figure 15:** **HEK293PAL cells transfected with eGFP shRNAs in darkness indicate different eGFP knockdown efficiencies.** Number of eGFP positive cells after transfection with the indicated shRNA. Data is also shown in **Fig. 2 c**,**f**,**i**. This figure magnifies the lower part of the y-axis to clarify eGFP knockdown differences of cells incubated in darkness. Shown are normalized values to SHA in darkness. N = at least three independent experiments performed in duplicates. dark grey bars: cells incubated in darkness. Values are means ± s.d.

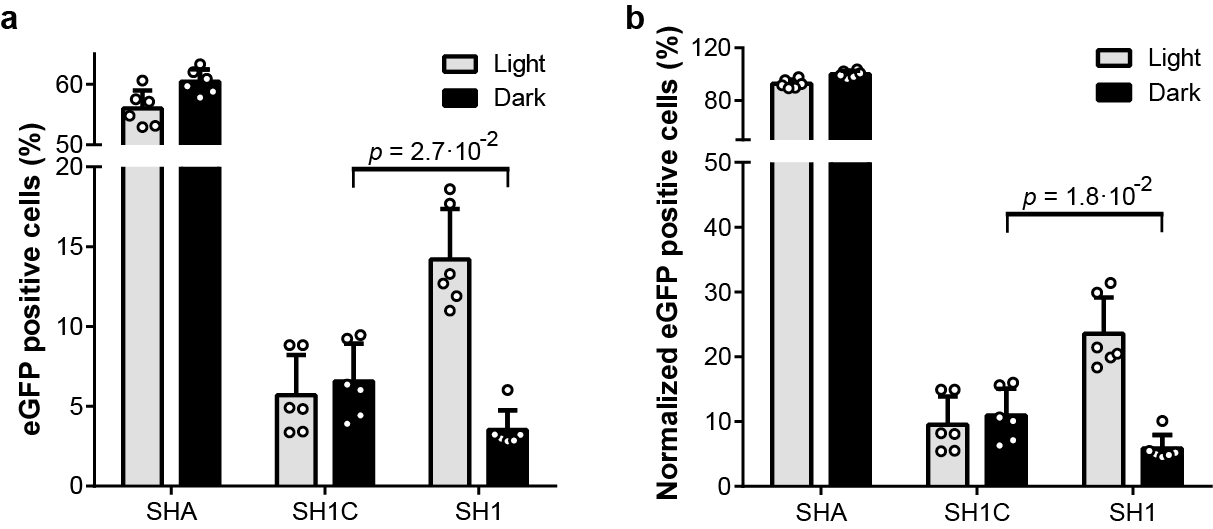

**Supporting Figure 16: Effect of the aptamer as apical loop domain on eGFP expression. a**, eGFP positive cells after transfection with the indicated shRNA variants. **b**, Normalized eGFP positive cells after transfection with the indicated shRNA. **b**, Values are normalized to SHA in darkness. **a**,**b**, eGFP positive cells were identified as indicated in **Supporting Fig. 6b**. **a**,**b**, Experiment was performed in duplicates and three independent replicates. **a**,**b**, Two-sided Mann-Whitney *U* test was used for statistical analysis as an unpaired observation was assumed. Grey bars: light conditions, black bars: dark conditions. Values are means ± s.d.

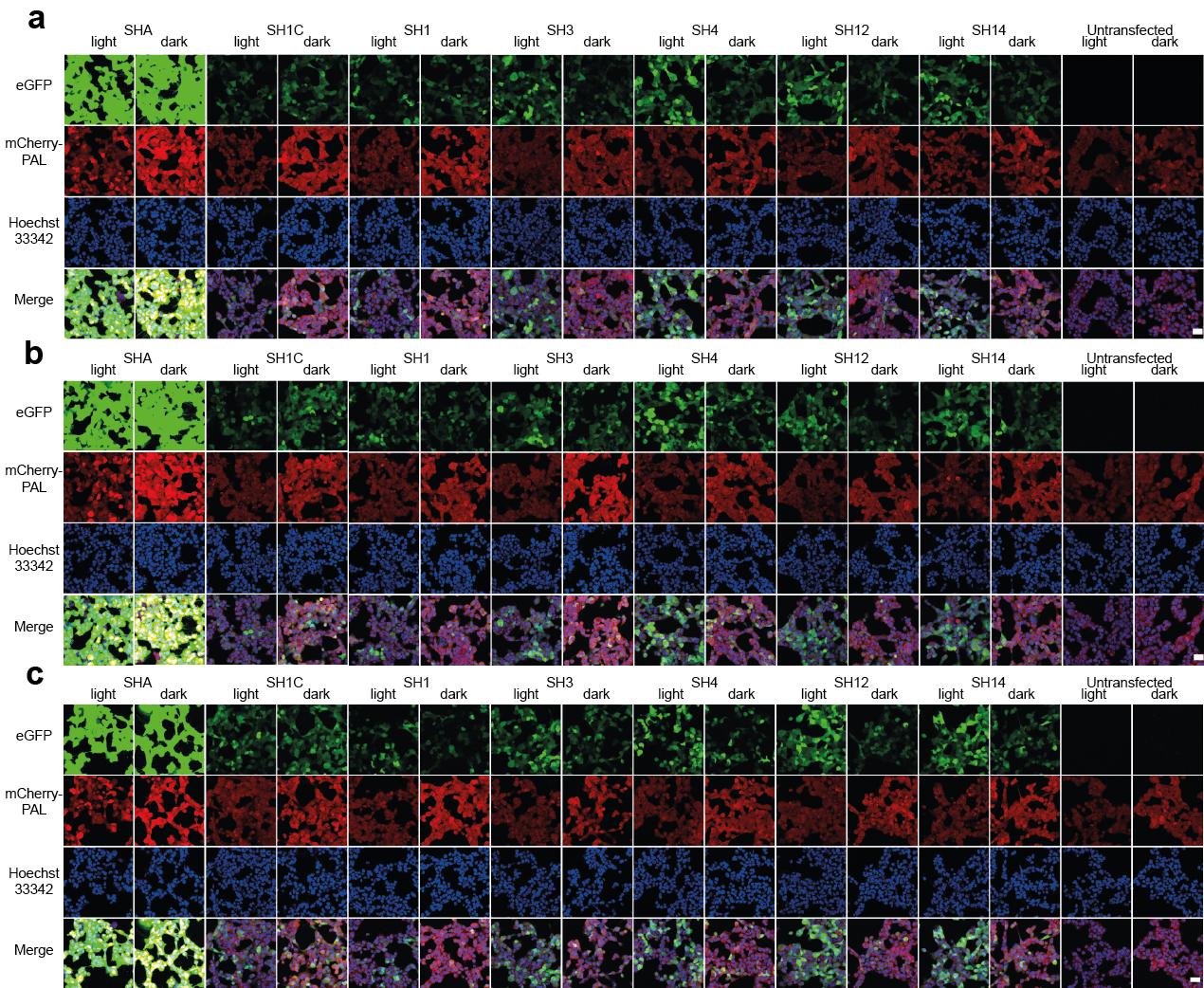

**Supporting Figure 17. Fluorescence microscopy pictures of HEK293PAL cells transfected with eGFP shRNAs indicate light-dependent eGFP expression.** Fluorescence microscopy images of HEK293PAL cells transfected with the indicated shRNAs. N = three independent experiments performed in duplicates. Cells were incubated under the indicated light conditions. **a**, **b**, **c**, representatives of first, second and third repetition, respectively. Data corresponds to **Fig. 2o**, where parts of **a** are shown. Scale bar: 40 μm.

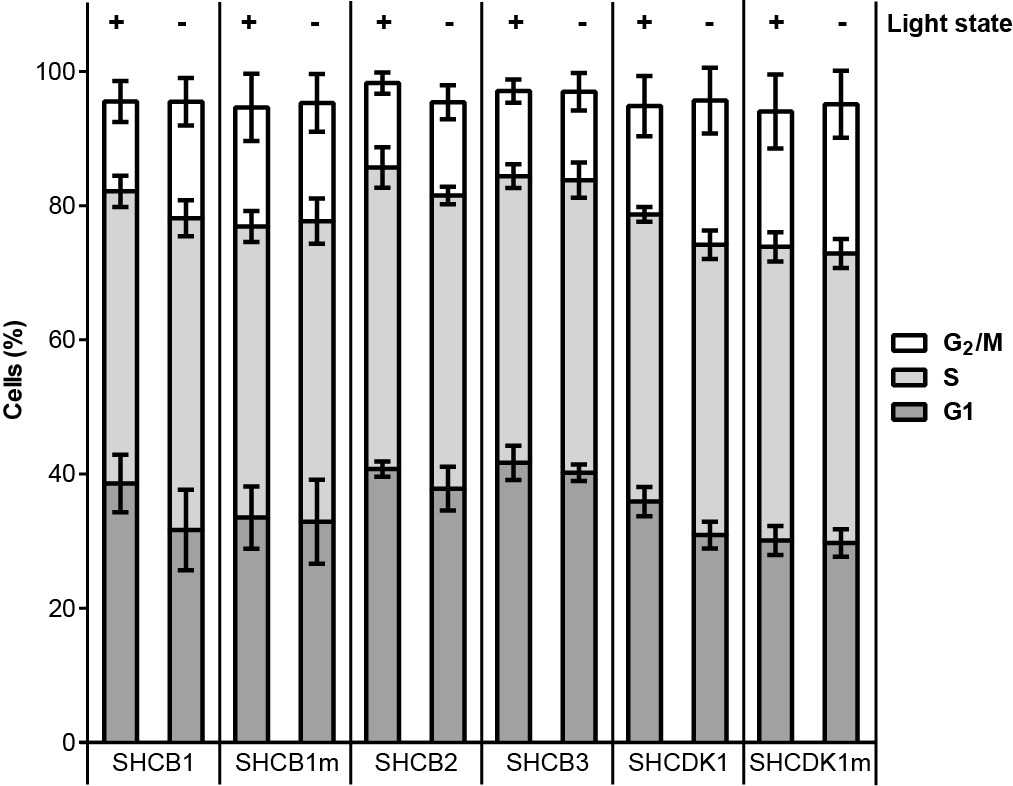

**Supporting Figure 18:** **Cell cycle phase distribution of cells transfected with shRNAs targeting cyclin B1 and CDK1.** Percentages of HEK293PAL cells in G_1_, S and G_2_/M phase of cells also shown in **Fig. 3b** and **d** after transfection with the indicated shRNAs. Identity of SHCB1, SHCB1m, SHCDK1 and SHCDK1m was blinded and double-blinded in one experiment, each. N = at least three independent experiments performed in duplicates. Experiments were performed under the indicated light conditions. Values are means ± s.d.

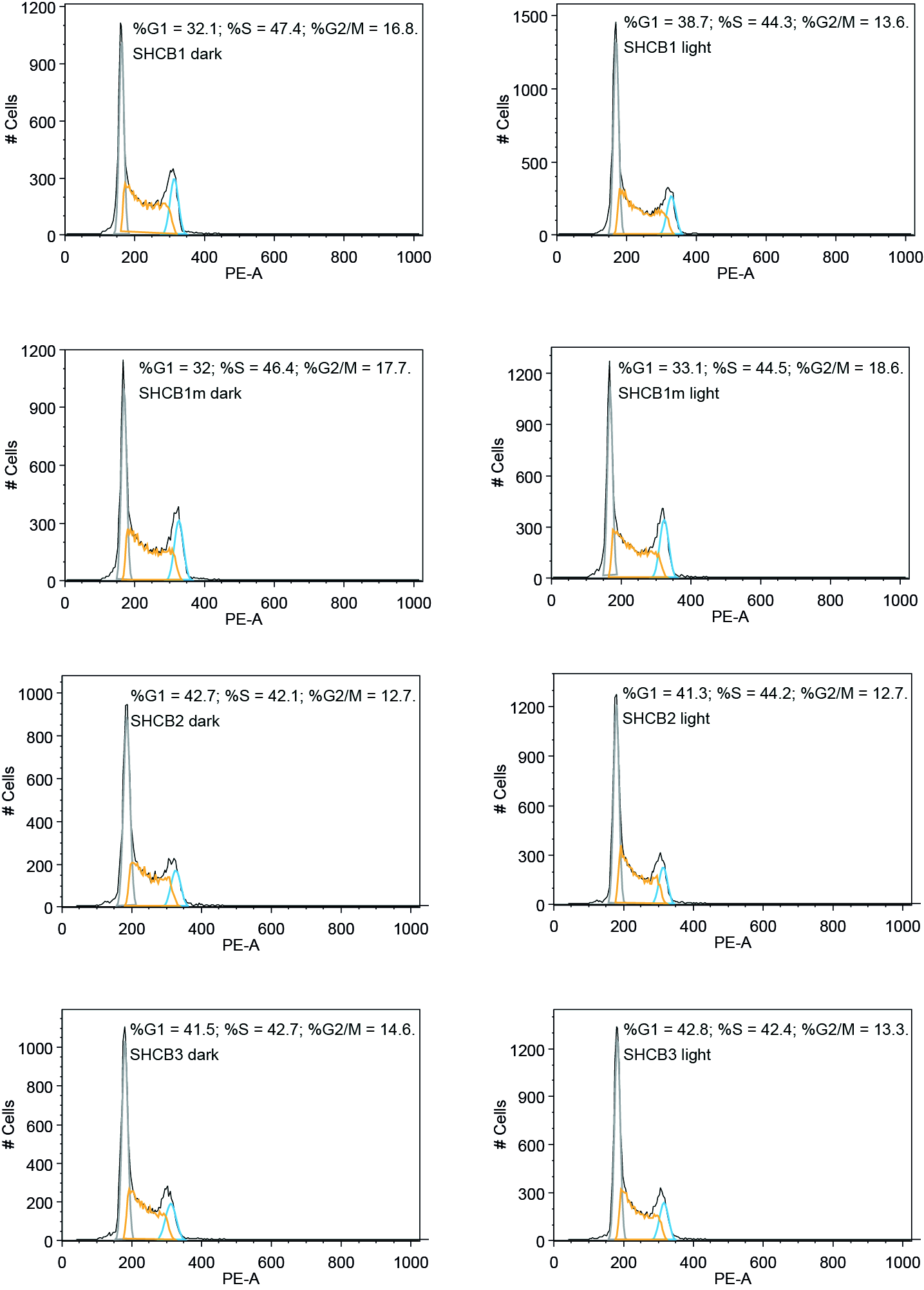

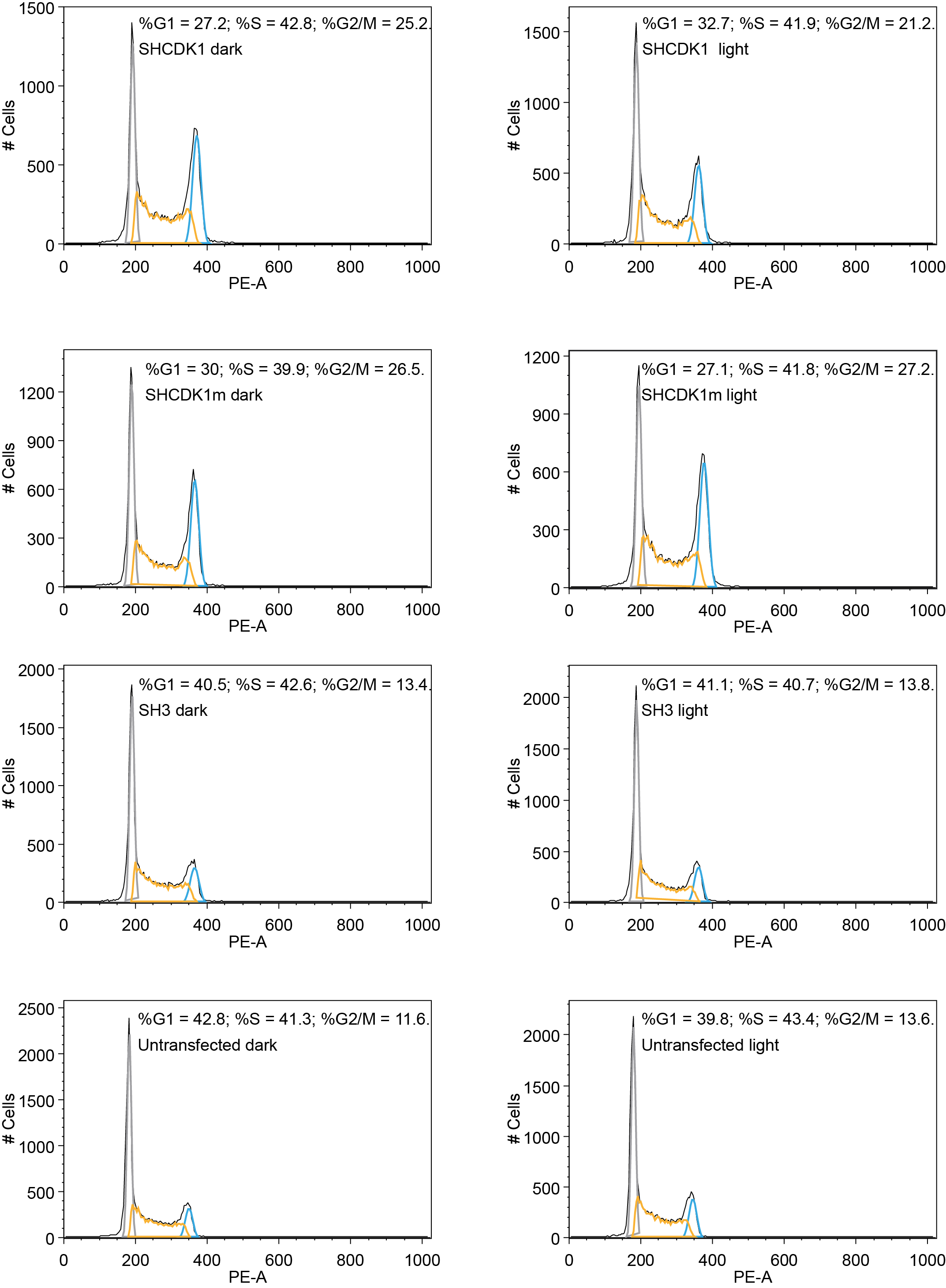

**Supporting figure 19: Cell cycle phase distribution is optoribogenetically controllable.** Representative crude flow cytometry diagrams of HEK293PAL cells transfected with the indicated shRNA variants or non-treated. Cell cycle phase percentages were calculated using Watson pragmatic algorithm. Grey: G_1_ phase, orange: S phase, blue: G_2_/M phase. N = at least three independent experiments performed in duplicates.

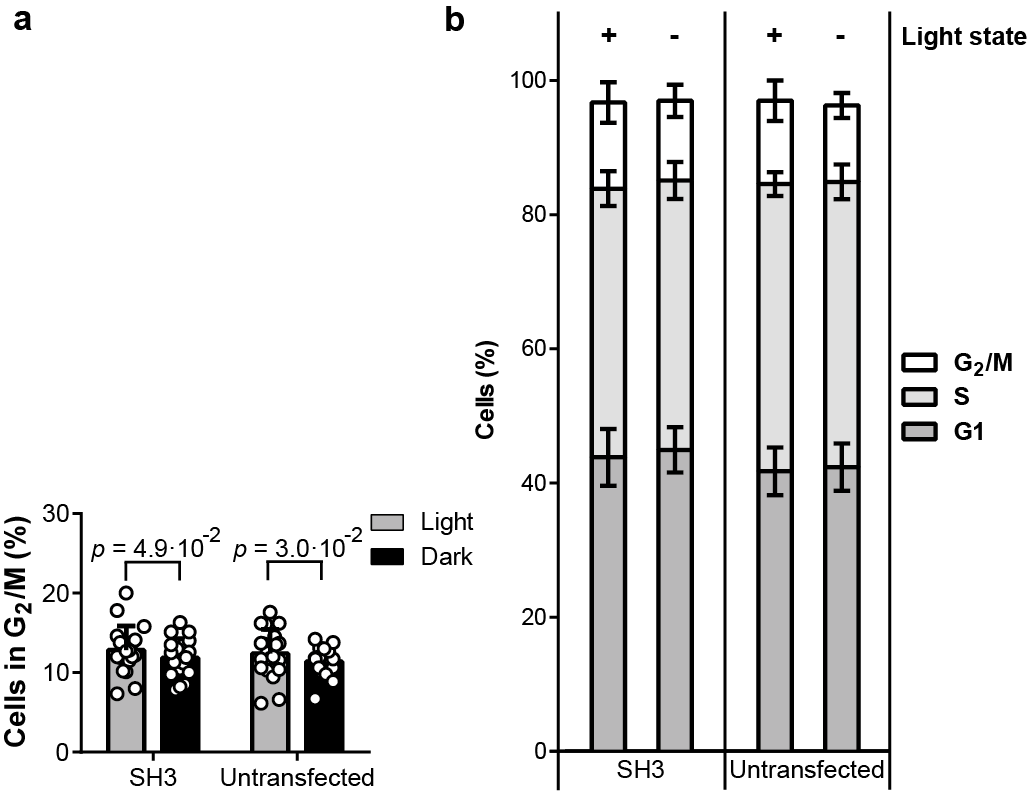

**Supporting Figure 20: Blue light slightly influences cell cycle distribution. a**, Percentages of HEK293PAL cells in G_2_/M phase after transfection with SH3 or no treatment. Grey bars: cells incubated under light conditions, black bars: cells incubated in darkness. **a**, Wilcoxon two-sided signed-rank test was used for statistical analysis as a paired observation was assumed. **b**, Percentages of HEK293PAL cells in G_1_, S and G_2_/M phase of cells shown in (**a**). **b**, Experiment was performed under the indicated light conditions. **a**,**b**, N = ten independent experiments performed in duplicates. **a**,**b**, Identity of SH3 was blinded and double-blinded in one experiment, each.. Values are means ± s.d.

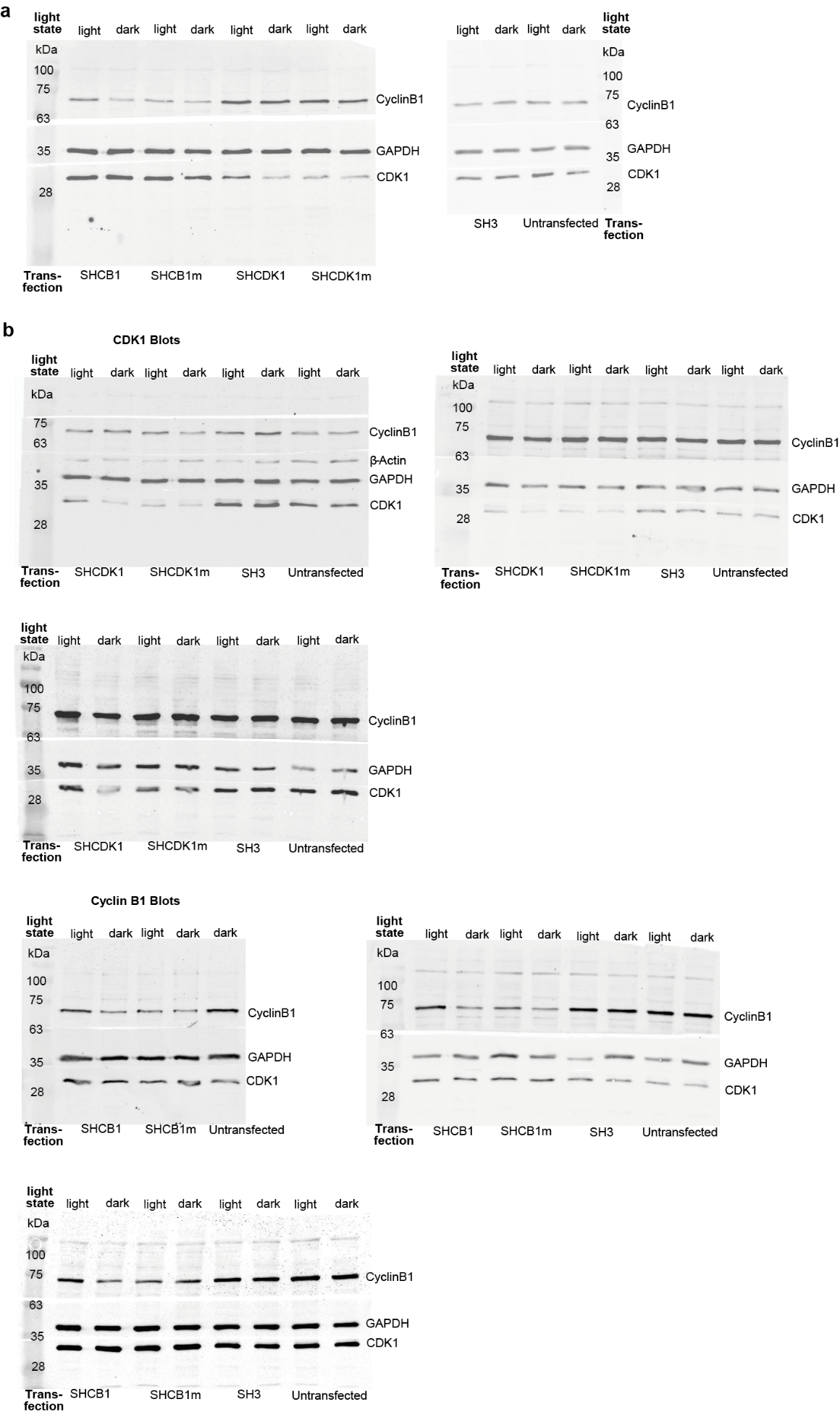

**Supporting Figure 21: Optoribogenetic control of target protein expression**. Full range Western Blots showing cyclin B1, CDK1 and GAPDH protein expression after transfection with the indicated shRNAs and the indicated light state. Parts of (**a**) were shown in **fig. 3e** (**a**). Western blots shown in (**b**) were used for quantification of cyclin B1 and CDK1 protein levels using pixel densitometry shown in **Fig. 3 f,g** (**b**). **b**, N = Three independent experiments.

**Supporting Table 1.** List of *pre*-miR21 and shRNA constructs used in this study. Blue: Aptamer sequence, red: Point Mutations, orange: Hinge region nucleotides, brown (bold): control loop sequence.

| **Name** | **RNA sequence (5’-3’)** |
| --- | --- |
| 53 | GGGAGGACGAUGCGGCCCGTACAGCAGCGATGCGGGTCGCGTGTCCACCCCGGCTCAGACGACUCGCUGAGGAUCCGAGA |
| SHA | GUAGCUUAUCAGACUGAUGUUGACGGUACAGCAGCGAUGCCGCAACACCAGUCGAUGGGCUGUUU |
| SHB | GAAGGCAAGCUGACCCUGAAGUUCGGUACAGCAGCGAUGCCGAAGGCAAGCUGACCCUGAAGUUU |
| SHC | GUAGCUUAUCAGACUGAUGUUGACGGUACAGCACCGAUGCCGCAACACCAGUCGAUGGGCUGUUU |
| SHD | GAAGGCAAGCUGACCCUGAAGUUCGGUACAGCACCGAUGCCGAAGGCAAGCUGACCCUGAAGUUU |
| SH1C | GCAAGCUGACCCUGAAGUUCA**UUCAAGAGA**UGAACUUCAGGGUCAGCUUGCUU |
| SH1 | GCAAGCUGACCCUGAAGUUCAUCGGUACAGCAGCGAUGCCGAUGAACUUCAGGGUCAGCUUGCUU |
| SH2 | GCACAAGCUGGAGUACAACUACGGUACAGCAGCGAUGCCGUAGUUGUACUCCAGCUUGUGCUU |
| SH3 | GCAAGCUGACCCUGAAGUUCAUACGGUACAGCAGCGAUGCCGAUGAACUUCAGGGUCAGCUUGCUU |
| SH4 | GCAAGCUGACCCUGAAGUUCAUUCGGUACAGCAGCGAUGCCGAUGAACUUCAGGGUCAGCUUGCUU |
| SH5 | GCAAGCUGACCCUGAAGUUCAUGCGGUACAGCAGCGAUGCCGAUGAACUUCAGGGUCAGCUUGCUU |
| SH6 | GCAAGCUGACCCUGAAGUUCAUCCGGUACAGCAGCGAUGCCGAUGAACUUCAGGGUCAGCUUGCUU |
| SH7 | GCAAGCUGACCCUGAAGUUCAUCGGUACAGCAGCGAUGCCGAAUGAACUUCAGGGUCAGCUUGCUU |
| SH8 | GCAAGCUGACCCUGAAGUUCAUCGGUACAGCAGCGAUGCCGUAUGAACUUCAGGGUCAGCUUGCUU |
| SH9 | GCAAGCUGACCCUGAAGUUCAUCGGUACAGCAGCGAUGCCGGAUGAACUUCAGGGUCAGCUUGCUU |
| SH10 | GCAAGCUGACCCUGAAGUUCAUCGGUACAGCAGCGAUGCCGCAUGAACUUCAGGGUCAGCUUGCUU |
| SH11 | GCACAAGCUGGAGUACAACUAACGGUACAGCAGCGAUGCCGUAGUUGUACUCCAGCUUGUGCUU |
| SH12 | GCACAAGCUGGAGUACAACUAUCGGUACAGCAGCGAUGCCGUAGUUGUACUCCAGCUUGUGCUU |
| SH13 | GCACAAGCUGGAGUACAACUACGGUACAGCAGCGAUGCCGAUAGUUGUACUCCAGCUUGUGCUU |
| SH14 | GCACAAGCUGGAGUACAACUACGGUACAGCAGCGAUGCCGUUAGUUGUACUCCAGCUUGUGCUU |
| SH15 | GCACAAGCUGGAGUACAACUACGGUACAGCAGCGAUGCCGGUAGUUGUACUCCAGCUUGUGCUU |
| SH16 | GCACAAGCUGGAGUACAACUACGGUACAGCACCGAUGCCGUAGUUGUACUCCAGCUUGUGCUU |
| SHCB1 | GACACCAACUCUACAAUAUUAGUUAACGGUACAGCAGCGAUGCCGUAGCUAAUGUUGUAGAGUUGGUGUCUU |
| SHCB1m | GACACCAACUCUACAAUAUUAGUUAACGGUACAGCACCGAUGCCGUAGCUAAUGUUGUAGAGUUGGUGUCUU |
| SHCB2 | GACACCAACUCUACAAUAUUAGUUAUCGGUACAGCAGCGAUGCCGUAGCUAAUGUUGUAGAGUUGGUGUCUU |
| SHCB3 | GACACCAACUCUACAAUAUUAGUUACGGUACAGCAGCGAUGCCGUUAGCUAAUGUUGUAGAGUUGGUGUCUU |
| SHCDK1 | GUGGAAUCUUUACAGGACUAUCACGGUACAGCAGCGAUGCCGGAUAGUCCUGUAAAGAUUCCACUU |
| SHCDK1m | GUGGAAUCUUUACAGGACUAUCACGGUACAGCACCGAUGCCGGAUAGUCCUGUAAAGAUUCCACUU |

**Supporting Table 2.** Sequencing of mature miR21-5p clones from SHA transfection indicates altered 3’-isomer formation compared to natural *pre*-miR21. Percentage was calculated from the number of clones obtained for the respective 3’-isomer. Blue: Nucleotides that code for the PAL-aptamer.

| **Name** | **# Clones** | **Percentage (%)** | **RNA sequence** |
| --- | --- | --- | --- |
| Mature miR21-5p | 0 | 0 | UAGCUUAUCAGACUGAUGUUGA |
| Mature miR21-5p – 1 nts | 2 | 17 | UAGCUUAUCAGACUGAUGUUG |
| Mature miR21-5p + 1 nts | 2 | 17 | UAGCUUAUCAGACUGAUGUUGA**C** |
| Mature miR21-5p + 2 nts | 8 | 67 | UAGCUUAUCAGACUGAUGUUGA**CG** |
